# VECTR-Clasp: An open machine learning and vector-based framework for objective quantification of hind-limb clasping and motor dysfunction in *Cdkl5*-deficient mice

**DOI:** 10.64898/2026.02.25.707643

**Authors:** Jordan Higgins, Samuel Egan, Keelin Harrison, Bilal El-Mansoury, David C. Henshall, Omar Mamad

**Affiliations:** Department of Physiology C Medical Physics, RCSI University of Medicine C Health Sciences, Dublin, D02 YN77, Ireland; FutureNeuro Research Ireland Centre for Translational Brain Science, RCSI University of Medicine C Health Sciences, Dublin, D02 YN77, Ireland; Department of Psychiatry, RCSI University of Medicine C Health Sciences, Dublin, D02 YN77, Ireland

**Keywords:** Hind-Limb Clasping, CDKL5, Computational Behaviour, Machine Learning, Motor dysfunction

## Abstract

Quantitative assessment of motor behaviour in rodents is central to the study of neurological disease, yet it remains constrained by manual categorical scoring systems that limit sensitivity, reproducibility, and the ability to detect subtle phenotypes. The hind-limb clasping assay, a standard test of motor dysfunction across many mouse models, is typically scored on an observer-defined categorical scale that may overlook meaningful movement features. We developed VECTR-Clasp, an open vector-based geometric framework that transforms standard pose-estimation output into continuous, body-relative kinematic measures. Combining DeepLabCut for markerless pose estimation with SimBA for automated clasping classification, our pipeline first reproduces conventional clasping detection at a level of agreement approaching that between trained human raters, and then extracts continuous geometric descriptors, including head directionality, total distance travelled, and swing count, that are inaccessible to categorical scoring. Applied to a mouse model of CDKL5 deficiency disorder, a rare neurodevelopmental condition in which motor impairment is a core clinical feature, this approach revealed previously uncharacterised motor microphenotypes: affected animals showed more constrained head direction, reduced overall movement, and fewer swings than wildtype controls. Critically, these differences were also present in affected animals that displayed no overt clasping. Together, these findings demonstrate that continuous geometric analysis of pose-estimation data uncovers motor phenotypes beyond the resolution of categorical scales, providing a more sensitive and reproducible framework for quantifying motor dysfunction. As it operates on standard pose-estimation output, the approach extends readily to other behavioural assays, disease models, and the evaluation of therapeutic interventions.

**Author Summary:** Many neurological disorders impair movement, and researchers frequently study these deficits in mouse models. A widely used test lifts a mouse by the tail and records whether it retracts its hind limbs towards its body, a reflex associated with motor dysfunction. Traditionally, an observer watches the animal and assigns a categorical score. This approach depends on human judgement, varies between raters, and can miss subtle movement differences.

We developed VECTR-Clasp, an open-source pipeline that validates machine-learning pose tracking to automate traditional clasping assessment and extends current methods using vector-based geometry. This quantifies the direction and extent of head and body movements, providing additional motor readouts. In a mouse model of CDKL5 deficiency disorder, a rare paediatric neurological disorder, VECTR-Clasp matched human scoring while revealing kinematic differences missed by standard assessment, including in animals appearing unaffected. This objective, sensitive framework could detect subtle motor changes and treatment responses across neurological diseases.

## Introduction

Assessment of rodent movement and cognition remains a cornerstone of neuroscience research. Behaviour can be understood as an observable output of the summation of neural computation in respect to both intrinsic and extrinsic stimuli, providing an insight into neurological status in a readily accessible and non-invasive manner (1). Rodent-based behavioural assays have historically provided critical insights into neural functioning such as in the case of place cell discovery (2), which would not have been possible without a behavioural assay aspect. Behavioural assays also play a crucial role in the evaluation of drug development, particularly in diseases where comorbidities are present, as in epilepsy (3), or when the symptoms are largely psychological, as in depression (4).

Many neurological disorders have a motor-related impairment. A classic motor impairment assessment in rodents is the hind-limb clasping assay. The assay involves lifting the mouse by its tail and measuring the duration of the retraction of hind limbs towards the torso of the mouse (5), which occurs in place of the expected placing instinct of mice. The hind-limb clasping assay has been used in many rodent models of neurological disease including Rett Syndrome (6), Charcot-Marie-Tooth (7), Parkinson’s Disease (8), Multiple Sclerosis (9), Amyotrophic Lateral Sclerosis (10), Cerebellar Ataxia (11), Alzheimer’s Disease (12), and Huntington’s Disease (13). There have been many brain regions implicated with this reflex including the cerebellum, basal ganglia, and neocortex, and it has been hypothesised that the mechanistic pathways involve the cerebello-cortico-reticular and cortico-striato-pallido-reticular pathways, which may be due to changes in noradrenaline and serotonin transmission (5).

Cyclin-dependent kinase-like 5 (CDKL5) deficiency disorder (CDD) is a rare neurodevelopmental disorder with an estimated incidence of 2.36 per 100 000 births (14). CDD is caused by mutations in the X chromosome-linked *CDKL5* gene and presents with the core symptoms of early-onset epilepsy, autism spectrum disorder-like traits and significant motor impairments (14). *Cdkl5*-deficient mice similarly recapitulate abnormal motor functioning and impairment seen in patients (15,16), as reflected by rodent-specific phenotypes including the hind-limb clasping reflex abnormality where a mouse displays a rigid posture with retracted hind-limbs. This appears to be a reversible phenotype and studies testing treatments for CDD have used it as therapeutic evaluation metric in mice (17).

Currently, hind-limb clasping is assessed using a semi-quantitative visual assessment and 5-point scoring system (11). This suffers the limitation of inter-user error, limited range of analyses, and need for repeated tester training. Thus, there is a need for fully objective and reproducible analysis methods that may expand the range of assessments and be driven by continuous kinematics rather than categorical scoring, which may be missing subtle motor phenotypes (18). Machine learning is now enabling such analyses, with large data driven frameworks in computational neuroethology and behavioural neuroscience applying tools such as DeepLabCut (19) and SimBA (20). These allow for increased objectivity and readouts from traditional behavioural assays such as beam walking (21), gait analysis (22), forced-swim test (23), novel object recognition (24), and social-based assays (25). Notably, they have allowed for traditionally difficult to track subtle phenotypes to be measured both longitudinally (26) and also assess the effects of drug treatments (27). In this study, we describe an approach using DeepLabCut and SimBA based pipelines to automate the quantification of clasping duration to capture this behaviour.

Advances in computational neuroethology have revealed that rodent behaviour is organised into reproducible, sub-second building blocks (28). Wiltschko et al. demonstrated that spontaneous mouse movements can be decomposed into distinct behavioural syllables lasting only a few hundred milliseconds and were associated with discrete neural activity patterns in motor and striatal circuits (28). Extending this, Markowitz et al. showed that the dorsolateral striatum encodes the identity and sequential organization of these sub-second motifs, supporting moment-to-moment action selection (29). Together, these studies established that fine-grained behavioural structure is not noise but a functional unit of neural computation.

Building on this conceptual framework, we developed a vector-based geometry approach that derives continuous kinematic measures of hind-limb clasping, such as head directionality and movement, directly from pose data. Unlike supervised classifiers, these measures are defined a priori, require no training labels, and remain computable for a single animal, providing a physically grounded description of movement rather than a learned discrimination score. We use the term ’microphenotype’ to refer to continuous kinematic features detectable by quantitative measurement but below the resolution of categorical scoring scales. Vector-based geometry has been successfully implemented in behavioural analysis before in the context of an open arena to quantify behavioural syllables (30) and to assess mobility parameters after injuries in rodents (31,32), but no attempts have been made for this specific assay. Despite its widespread use, the hind-limb clasping assay has never been analysed with continuous kinematic methods, leaving open the question of whether categorical scoring overlooks biologically meaningful motor features. We hypothesised that a vector-based geometric framework applied to pose estimation data would reveal continuous motor microphenotypes in *Cdkl5*-deficient mice that are undetectable by standard categorical scoring. Here we show that this approach achieves two things: it automates a traditionally subjective assay at close to the agreement between trained human raters, and it yields continuous, single-animal motor descriptors that expose genotype differences invisible to the categorical clasping scale, including in animals that show no overt clasping.

## Results

### Pose Estimation s Behavioural Classifier

We first conducted the hind-limb clasping assay as shown in Fig 1 to begin our data collection. To obtain pose estimation data for downstream analysis, DeepLabCut was employed. Here, we trained the pose estimation model on 15 one-minute videos from 15 mice, consisting of 5 wildtype, 6 knockout and 4 heterozygous mice. DeepLabCut accurately detected body parts across videos and achieved high confidence after training. The model achieved a Training/validation loss ≈ 0.002, a Root Mean Square Error (RMSE): 7.47 px, a RMSE at p-cutoff 0.6: 3.16 px, Mean Average Precision (mAP): 96.12%, and a Mean Average Recall (mAR): 97.10%. The best snapshot was chosen at epoch 145 (out of 200) based on training plateau and subsequent evaluation metrics for each following epoch performing worse or having negligible differences. Representative pose estimation plots obtained of wildtype and knockout are shown in Fig 2A.

**Fig 1.**
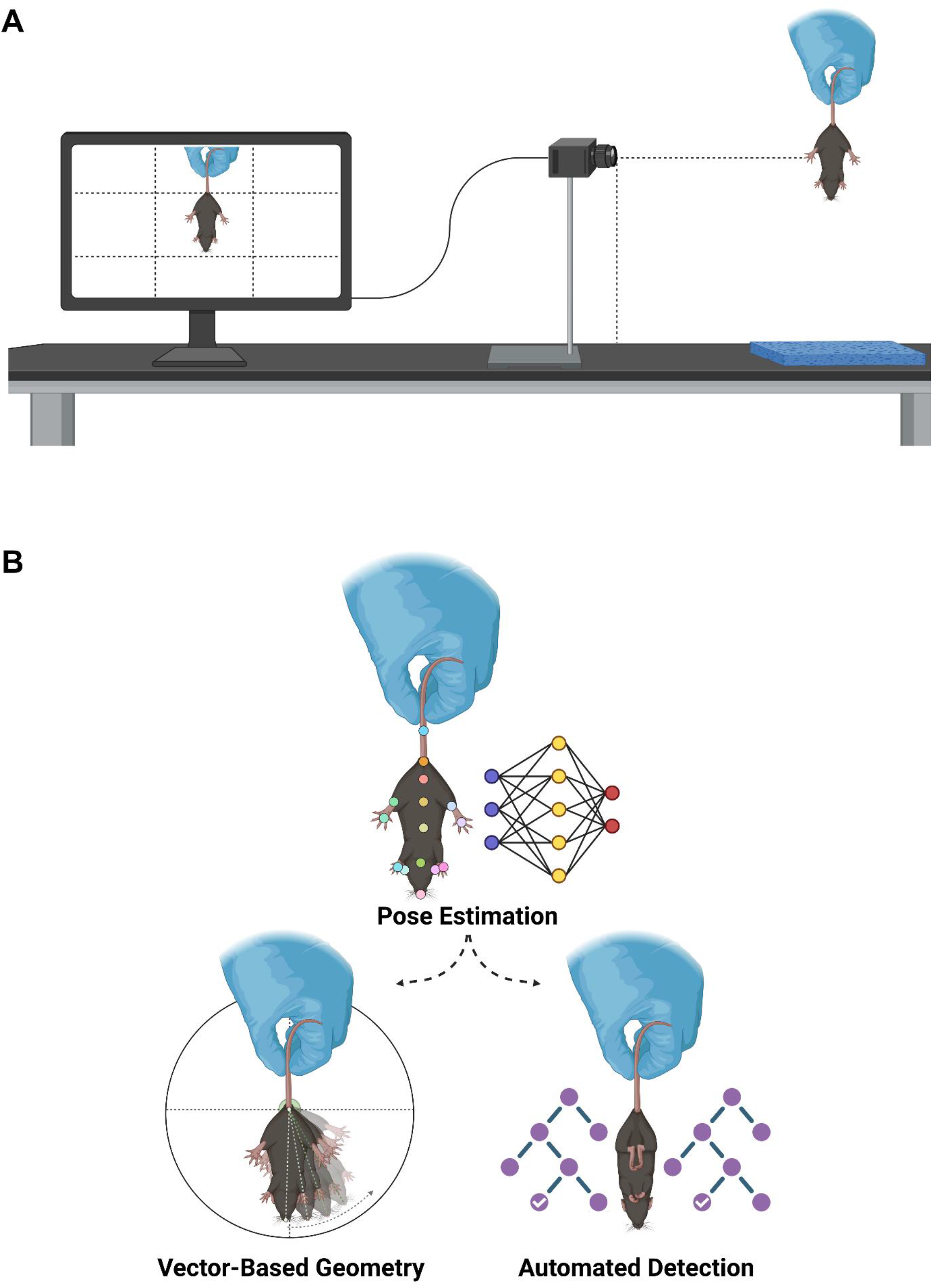
Automated and geometric Analysis of Hind-Limb Clasping Behaviour Using Deep Learning-Based Pose Estimation. (A) Experimental setup for video acquisition of hind-limb clasping behaviour. Dashed lines indicate a 30 cm distance. A soft blue pad was placed below the mouse to prevent injury in case of a fall. The computer screen was divided into nine sections, with the mouse positioned within the central third for standardised recording. (B) Schematic representation of the analysis pipeline combining deep learning-based pose estimation, automated behaviour classification, and geometric vector analysis. A total of 90 mice were used for pose-estimation with DeepLabCut (15 training). Behavioural classification in SimBA utilised 5 training and 70 held-out videos. While vector-based geometric analysis used only the 70 held-out videos.

**Fig 2.**
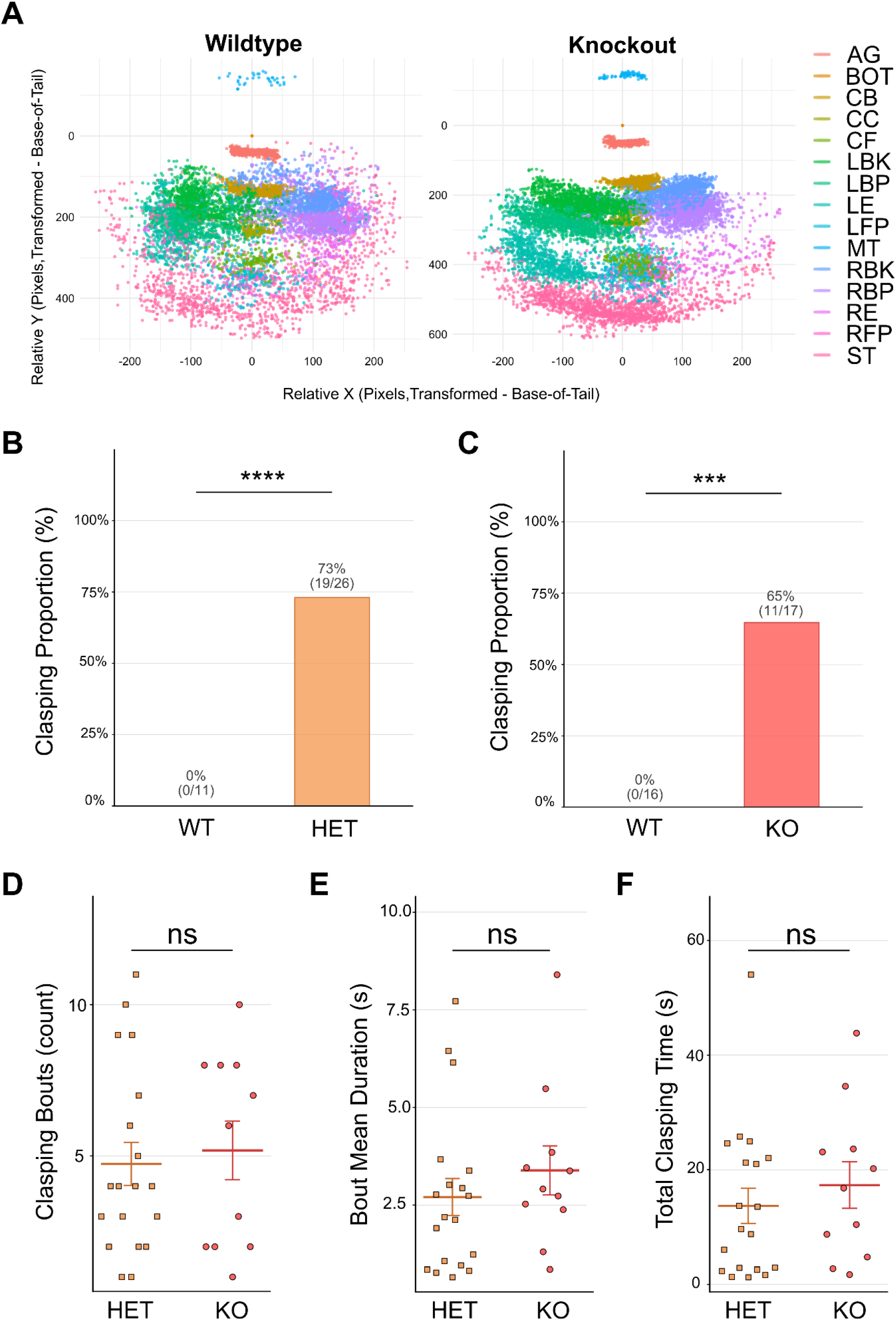
Quantitative and Temporal Analysis of Hind-limb Clasping Behaviour Across Genotypes. (A) Relative XY coordinates of tracked body parts in a wildtype and knockout mouse, aligned to the base of the tail (likelihood > 0.6). Each body part is color-coded as indicated in the legend. Data represent 60 seconds of recording (1800 frames). Knockout mice exhibit tighter clustering of body part coordinates, a more spatially constrained, less variable pattern of movement during tail suspension, whereas wildtype mice show greater spatial dispersion consistent with dynamic movement. Population-level quantification of these movement patterns is presented in Fig 5. AG: anogenital region, BOT: base of tail, CB: centre back, CC: centre centre, CF: centre front, LBK: left back knee, LBP: left back paw, LE: left ear, LFP: left front paw, MT: mid-tail, RBK: right back knee, RBP: right back paw, RE: right ear, RFP: right front paw, ST: snout tip. (B) Proportion of female wildtype (WT; n=11) and heterozygous (HET; n=26) animals exhibiting at least one clasping event, assessed by automated SimBA scoring. No WT females clasped; 73% (19/26) of HET females clasped (Fisher’s exact test, p < 0.0001). (C) Proportion of male wildtype (WT; n=16) and knockout (KO; n = 17) animals exhibiting clasping. No WT males clasped; 65% (11/17) of KO males clasped (p < 0.001). Error bars represent 95% confidence intervals. (D-F) Comparisons of clasping bout count (D), mean bout duration in seconds (E), and total clasping duration in seconds (F) between HET and KO animals that exhibited clasping. No significant differences were observed between groups on any measure (ns; Mann-Whitney U test, Bonferroni corrected). Individual data points shown; horizontal lines indicate median ± IQR.

To automate the clasping detection with a random forest classifier, SimBA was trained on 5 videos, consisting of 2 wildtype and 3 knockouts, after DeepLabCut analysis. The SimBA behavioural classifier model achieved high accuracy during training with a maximum F1.0 score of 0.959 and a maximum F0.5 score of 0.946 shown in Supplementary Fig 1. These values then informed the discrimination threshold values for clasping detection or rejection. The classifier extracted 2429 features from the geometric co-ordinates with the most important features, being relational geometric features indicating changing area between limbs (e.g. convex perimeter-based metrics), shown in Supplementary Fig 1. We chose the F0.5 model as the model to evaluate going forward, hereafter referred to as the SimBA model, and the proceeding results concern this chosen model.

As SMOTE oversampling might bias classifier performance, the model was evaluated on independent data. For SimBA model validation, we retained 70 hold-out videos consisting of 27 wildtype (11 female, 16 male), 26 heterozygous and 17 knockout mice (60 seconds per animal; 4,200 total scored seconds) shown in Table 1.

**Table 1:** Distribution of mice across analysis stages.

| Sex | Genotype | DeepLabCut Training | SimBA Training | SimBA and VECTR-Clasp Analysis | Genotype Classifier |
| --- | --- | --- | --- | --- | --- |
| Female | Wildtype | 5 | - | 11 | 11 |
|  | Heterozygous | 4 | - | 26 | 26 |
| Male | Wildtype | - | 2 | 16 | 19 |
|  | Knockout | 6 | 3 | 17 | 21 |

The number of mice contributing to each stage of the analysis pipeline is shown, stratified by sex and genotype. DeepLabCut pose estimation training used frames extracted from 15 videos (5 wildtype female, 4 heterozygous females, 6 knockout males). SimBA behavioural classifier training used 5 videos (2 wildtype male, 3 knockout male). SimBA and VECTR-Clasp geometric analysis was performed on 70 independent held-out videos not used at any prior stage of training or annotation (11 wildtype female, 26 heterozygous female, 16 wildtype male, 17 knockout male). The genotype classifier used the mice from both SimBA training and held-out evaluation plus the 2 mice removed from the SimBA-VECTR-Clasp analysis. A dash indicates that the genotype was not included at that stage. All mice were from the B6.129(FVB)-Cdkl5tm1.1Joez/J line.

A total of 11/17 knockouts, 19/26 heterozygous and 4/27 wildtype mice were detected to have clasping events. Following further investigation into these wildtype events, false positives were identified with only 1 bout detected and lasting less than 1 second. For proportion analysis, all mice with clasping duration above 1 second were considered as clasping to minimise false positives resulting from a spurious 1 bout-low duration detection. In this case, only the wildtype group was affected, and all four mice were deemed non-clasping.

To assess whether CDKL5 loss-of-function alters hind-limb motor behaviour, we analysed female heterozygous and male knockout mice alongside sex-matched wildtype controls, thus mimicking the clinically relevant populations. We first examined the binary clasping phenotype: no wildtype animals exhibited clasping behaviour during the 60-second trial (0/11 females; 0/16 males), whereas clasping occurred in 73% of heterozygous females (19/26; Fisher’s exact test, p[bonf] < 0.001) and 65% of knockout males (11/17; p[bonf] < 0.001; Fig 2B,C).

To determine whether clasping severity differed between genotypes, we compared per-clasping-animal characteristics between female heterozygous and male knockout animals that exhibited clasping. No significant differences were found between heterozygous (n=19; median bout duration=2.19 s [IQR: 1.01-3.20]) and knockout (n=11; median=2.91 s [IQR: 2.45-3.65]) animals in mean clasping bout duration (Mann-Whitney U, p[bonf] = 0.685), number of clasping bouts (heterozygous: median=4 [IQR: 2.5-6.5], knockout: median=6 [IQR: 2-8]; p[bonf] = 1.000), or total clasping duration (heterozygous: median = 9.63s [IQR: 2.70-21.63], knockout: median=16.80s [IQR: 6.75-23.37]; p[bonf] = 1.000).

### Human-Model Agreement

Next, we assessed the performance of the chosen SimBA model against human raters. Two human raters scored in a binary fashion per second, aiming to validate the model at a 1 second resolution. A clasping event was determined if the mouse had both hind limbs fully retracted. With this approach slight human-to-human variability was detected, although humans agreed mostly, highlighted by Fig 3A. Some edge cases, i.e. determining when a clasping event started or ended, being largely the source of disagreement, the same was true for the model after bringing the bout detection to the second epoch resolution as shown in Supplementary Fig 2.

**Fig 3.**
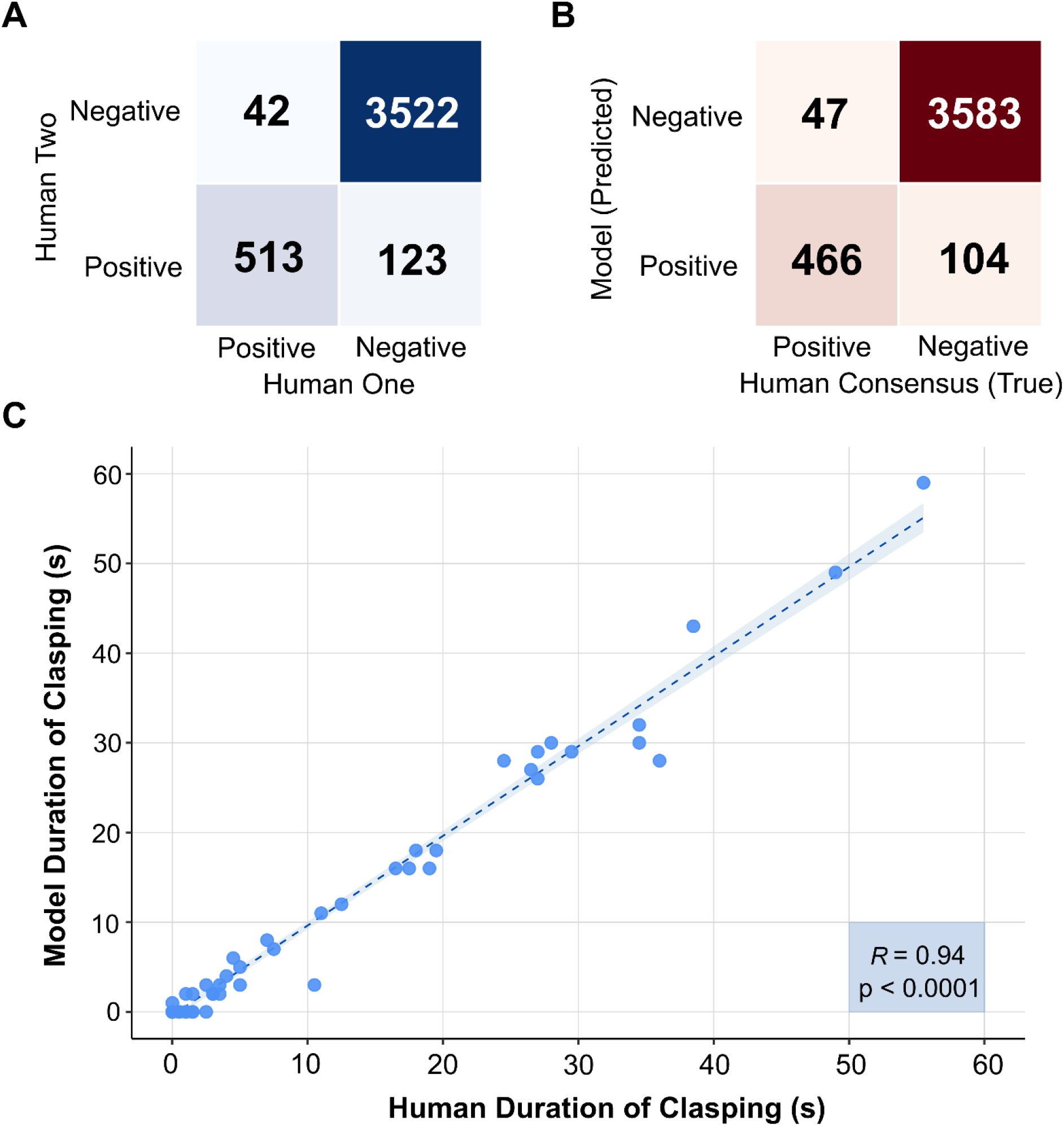
Agreement Between Human Raters and Automated Classification of Hind-limb Clasping Behaviour. (A) Confusion matrix comparing per-second clasping classifications between two independent human raters across all scored animals (4,200 total seconds). (B) Confusion matrix comparing per-second SimBA model predictions against the conservative human consensus label (seconds scored as clasping by both raters independently). The model achieved 96.4% overall agreement, with sensitivity 90.8% and specificity 97.2%. (C) Spearman correlation between total clasping duration scored by the model and total human-consensus clasping duration across individual animals. The substantial correlation (R = 0.94, p < 0.0001) confirms that the SimBA pipeline accurately captures individual variation in clasping duration.

Cohen’s Kappa (κ) values indicated substantial agreement across pairwise comparisons. The model and human intersection achieved a pooled κ = 0.84 [95% CI: 0.81-0.87]. Human-Human pooled κ = 0.84 [0.81-0.86], confirming strong inter-rater reliability and demonstrating that the model achieves similar agreement levels. The per-second epoch agreement was 96.4% (model-human) and 96.1% (human-human) across the full 4,200 scored seconds.

The model detected 466 true clasping and 3,583 true non-clasping epochs with 104 false positives and 47 false negatives, achieving high scores in accuracy (0.964), precision (0.818), and recall (0.908) as shown in Fig 3B. Following from this, a Spearman correlation was performed to assess how consistent the duration of scoring was between the model and the human intersection scoring. This achieved an R of 0.94 (Fig 3C), indicating very strong correlation between the two rating techniques (p < 0.0001). Of note, of all knockout and heterozygous mice which the model did not detect clasping, neither did the human consensus.

### Vector-Based Geometry

Having shown that traditional clasping scoring could be automated and accurately detected at 1 second resolution using DeepLabCut and SimBA, we next sought to further investigate this assay, generating continuous geometric measurements for deeper phenotyping of the movement patterns. To achieve this, we used the same 70 videos analysed by SimBA for geometric analysis. We first evaluated the percentage of frames which contained both the snout tip and the base of tail above a probability of 0.6 for our analysis and observed that wildtypes had an average of 77.5% frames, heterozygotes 81.5% frames and knockouts had 86.9% frames. Representative snout trajectories, probability densities heatmaps and polar histograms are shown in Fig 4 of wildtype, heterozygous and knockout mice.

**Fig 4.**
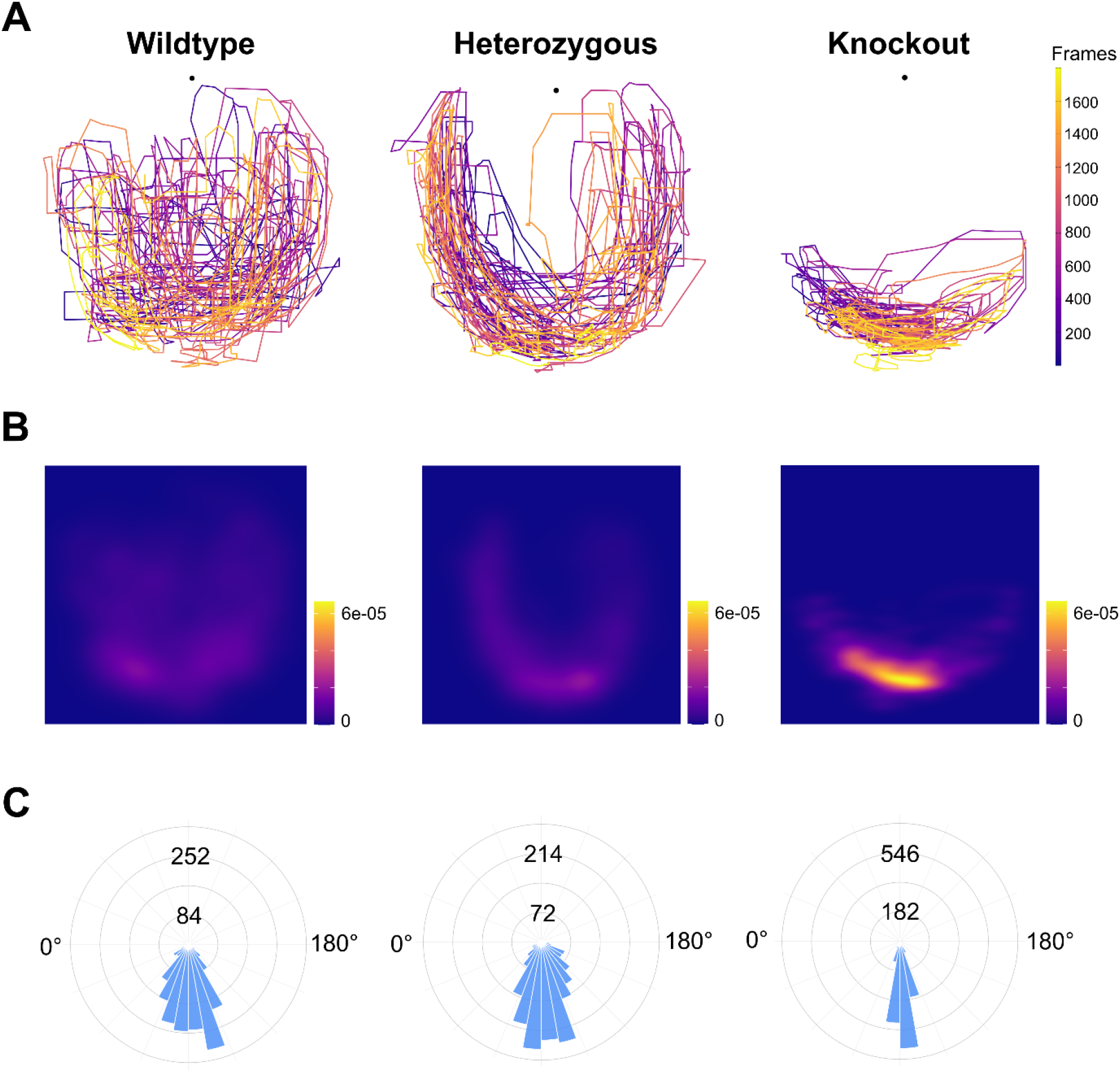
Spatial and Directional Analysis of Snout Movement During Tail Suspension. (A) Snout trajectory plots for representative wildtype, heterozygous, and knockout mice during the 60-second tail suspension trial. Trajectories are colour-coded by frame number (early = purple, late = yellow), illustrating the reduced spatial range of snout movement in affected animals. (B) Probability density heatmaps of snout position across all frames for the same representative animals, showing the spatial distribution of snout occupancy. Wildtype animals exhibit broad, bilateral snout coverage; heterozygous animals show an intermediate pattern; knockout animals display a markedly restricted and ventrally concentrated distribution. (C) Polar histograms of snout-to-tail-base angle across all frames. Wildtype animals show a wider angular spread, while knockout animals show a narrower, more stereotyped directional distribution. Numbers on the inner and outer rings indicate bin counts.

To determine whether sex independently influences mean resultant length, a linear model was fitted for wildtype animals only, with sex, age, and body size as predictors. Sex was a significant predictor of mean resultant length (p = 0.004), with wildtype males showing higher mean resultant length than wildtype females after covariate adjustment (estimated marginal means: 0.894 vs 0.817). Neither age (p = 0.133) nor a pixel-derived body size proxy, calculated as the ratio of tail-base-to-snout length to waistline width from the recording video, reached significance (p = 0.529). This proxy serves as an approximation of body size but is not equivalent to direct weight measurement, and the non-significant result should be interpreted accordingly. The model explained 35.7% of variance in mean resultant length (adjusted R² = 0.273), with sex accounting for the majority of explained variance. These findings indicate that males have an inherently higher baseline mean resultant length than females, independent of age and body composition, and should be considered when interpreting genotype-driven differences. Accordingly, subsequent analyses were stratified by sex comparing male wildtype and knockout animals, and female wildtype and heterozygous animals to accurately represent the clinically relevant populations.

Stratified analyses revealed distinct geometric patterns between groups (Fig 5A-D), with wildtype mice exhibiting greater variability in head direction and total distance compared to heterozygous and knockout mice. Head directionality and angular spread were quantified using mean resultant length and circular standard deviation (SD). In males, mean resultant length was significantly higher in knockout compared to wildtype animals (wildtype: mean = 0.886 ± 0.008 SEM, knockout: mean = 0.938 ± 0.007 SEM; p = 5.74 × 10⁻⁵, adjusted p = 5.74 × 10⁻⁴), with lower circular SD (wildtype: mean = 0.487 ± 0.019 rad, knockout: mean = 0.348 ± 0.023 rad; p = 5.92 × 10⁻⁵, adjusted p = 5.92 × 10⁻⁴), indicating a less dispersed angular distribution in knockouts. Comparable patterns were observed in females, with wildtype animals showing higher mean resultant length (wildtype: mean = 0.829 ± 0.018 SEM, heterozygous: mean = 0.899 ± 0.010 SEM; p = 0.004, adjusted p = 0.036) and lower circular SD (wildtype: mean = 0.606 ± 0.037 rad, heterozygous: mean = 0.444 ± 0.028 rad; p = 0.002, adjusted p = 0.020) than heterozygous animals.

**Fig 5.**
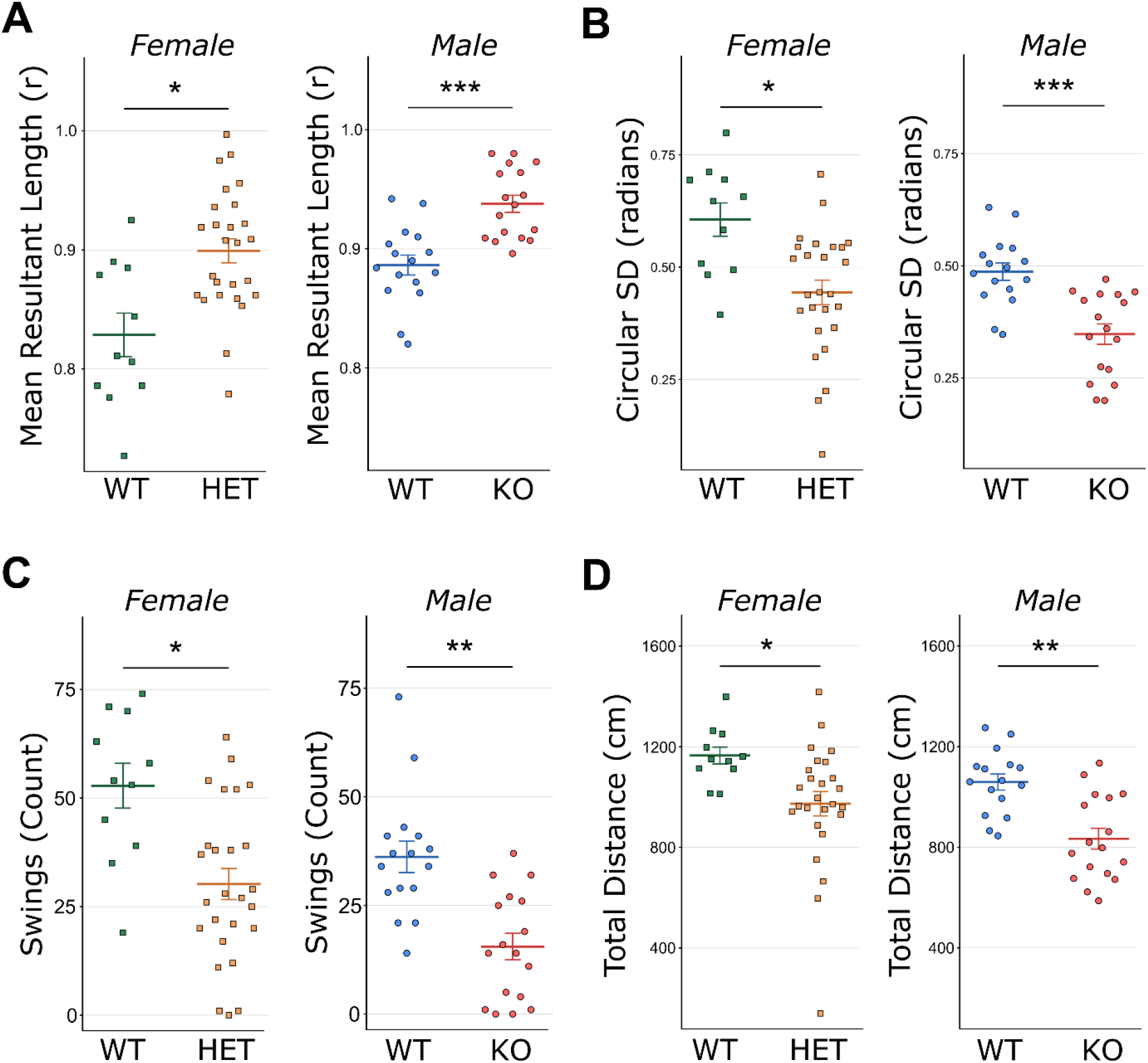
Geometric Quantification of Hind-limb Clasping Phenotypes. (A) Mean resultant length (MRL) for female WT vs HET (left) and male WT vs KO (right). Higher MRL reflects reduced angular variability (more stereotyped head direction). (B) Circular standard deviation (SD) in radians for female WT vs HET and male WT vs KO. Lower circular SD indicates more constrained directional movement. (C) Swing count for female WT vs HET and male WT vs KO, reflecting the number of lateral pendulum swings detected during the trial. (D) Total distance (cm) for female WT vs HET and male WT vs KO. Significant genotype differences were observed across all measures in both sexes. Significance symbols reflect Bonferroni-corrected p-values.

Wildtype mice also showed greater overall locomotion (Supplementary Video 1), which was reflected in significantly greater total distance compared to knockouts in both male (wildtype: mean = 1058.97 ± 31.74 cm, knockout: mean = 834.18 ± 41.32 cm; p = 1.65 × 10⁻⁴, adjusted p = 1.65 × 10⁻³) and female (wildtype: mean = 1165.01 ± 33.68 cm, heterozygous: mean = 973.38 ± 48.11 cm; p = 0.004, adjusted p = 0.040) comparisons.

As an exploratory measure, a swing threshold was defined as an exit ±45° from the tail suspension hanging position. Using this approach, swing count was significantly reduced in knockout animals relative to wildtype in males (wildtype: median = 35.5 [28.75, 41.00], knockout: median = 14.0 [4.00, 26.00]; p = 1.33 × 10⁻⁴, adjusted p = 1.33 × 10⁻³) and in females (wildtype: median = 54.0 [42.00, 66.50], heterozygous: median = 27.5 [20.00, 39.00]; p = 0.002, adjusted p = 0.017), with a small proportion of knockout animals recording zero swings. Together, these measures describe a more spatially constrained, less variable pattern of movement in *Cdkl5*-deficient mice during tail suspension including reduced snout displacement, more concentrated head direction, and fewer angular excursions from the hanging axis that is present even in animals without overt clasping (Supplementary Video 2).

### VECTR-Clasp Detects Geometric Phenotypes Independent of Overt Clasping

A key question for the interpretation of VECTR-Clasp measures is whether the observed genotype differences reflect the clasping phenotype itself, or an independent geometric signal present across the full range of phenotypic severity. To address this, affected animals were stratified post-hoc into those showing zero clasping duration (heterozygous: n=7; knockout: n=6) and those showing at least one clasping event (heterozygous: n =19; knockout: n =11), and VECTR-Clasp measures were compared across three groups per sex (Fig 6A-F). Previously identified wildtype mice with below 1 second clasping duration were set as 0 after manual confirmation. As mean resultant length and circular SD are analytically derived from one another (Equation 7) and provide equivalent directional information, Fig 6 displays mean resultant length as the primary directional measure; significant circular SD differences are reported here for completeness.

**Fig 6.**
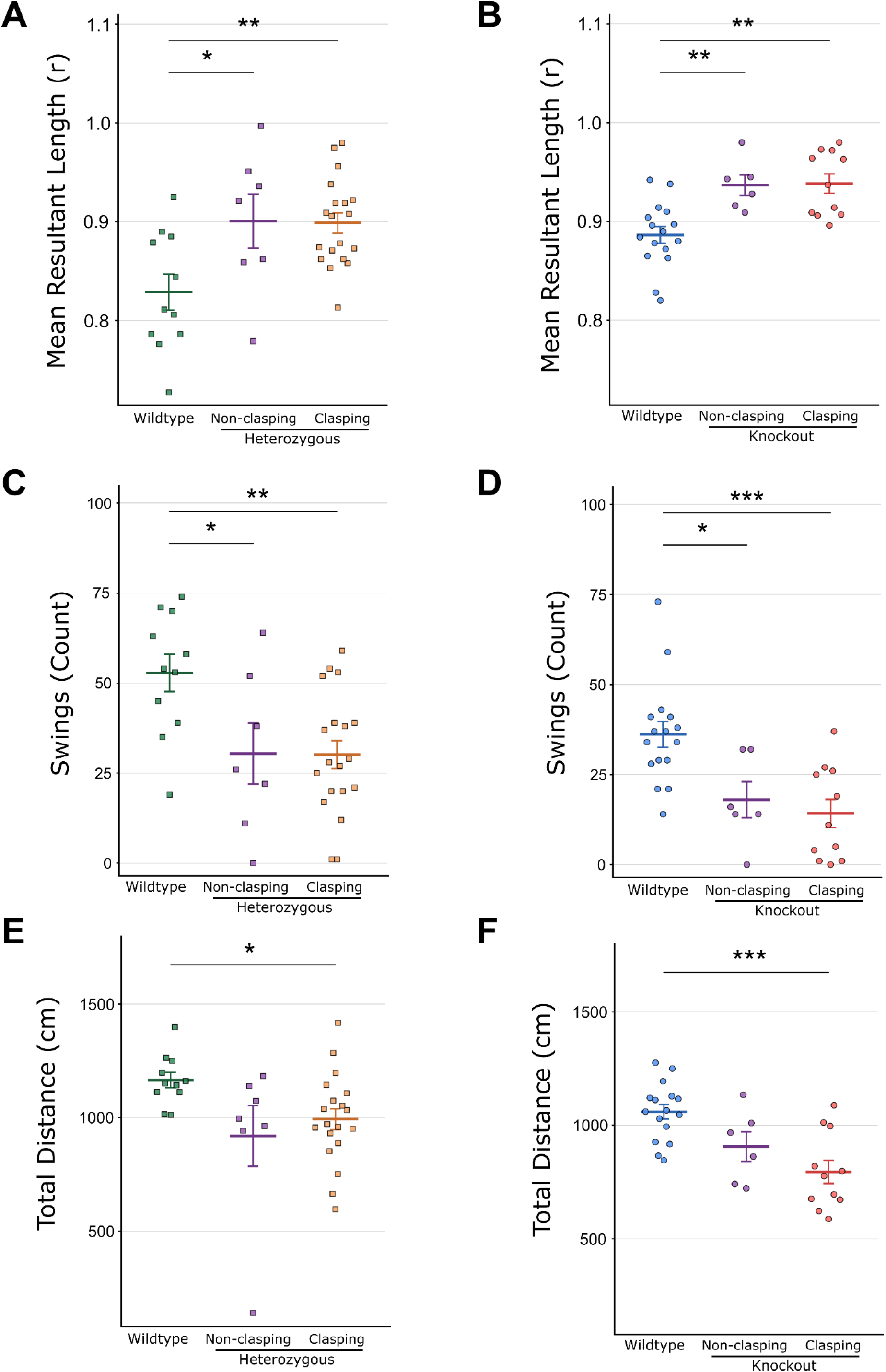
VECTR-Clasp kinematic phenotype is present in non-clasping mutant animals. Mutant animals were stratified post-hoc into those exhibiting zero clasping duration (non-clasping) and those exhibiting at least one clasping event (clasping), and compared against sex-matched wildtype controls. (A, B) Mean resultant length for heterozygous females (A) and knockout males (B). (C, D) Swing count for heterozygous females (C) and knockout males (D). (E, F) Total distance travelled for heterozygous females (E) and knockout males (F). As mean resultant length and circular SD are mathematically related (Equation 7), and therefore carry equivalent information, only mean resultant length is shown; circular SD results are reported in the Results text. Significance symbols reflect Bonferroni-corrected p-values.

In females, one-way ANOVA identified significant overall group differences for mean resultant length (F = 6.356, p = 0.005), circular SD (F = 5.493, p = 0.009), and swing count (F = 6.027, p = 0.006). A significant overall group difference was also observed for total distance (Kruskal-Wallis H = 8.358, p = 0.015). Post-hoc Tukey HSD tests revealed that non-clasping heterozygous animals - despite showing zero clasping duration by both human and automated scoring - had significantly higher mean resultant length (adjusted p = 0.028), lower circular SD (adjusted p = 0.027), and lower swing count (adjusted p = 0.039) relative to wildtype animals. Total distance in non-clasping heterozygous animals did not reach significance relative to wildtype (adjusted p = 0.097), though clasping heterozygous animals were significantly reduced compared to wildtype (adjusted p = 0.018). Critically, clasping and non-clasping heterozygous animals did not differ significantly on any measure (mean resultant length: adjusted p = 0.996; circular SD: adjusted p = 0.907; swing count: adjusted p = 0.999; total distance: adjusted p = 1.000), confirming that the geometric phenotype is not attributable to the presence or absence of overt clasping within the affected group.

In males, significant overall group differences were observed for VECTR-Clasp measures (mean resultant length: Kruskal-Wallis H = 14.413, p < 0.001; circular SD: F = 10.400, p < 0.001; total distance: F = 10.588, p < 0.001; swing count: F = 9.625, p < 0.001). Post-hoc comparisons confirmed that non-clasping knockout animals were significantly different from wildtype on mean resultant length (adjusted p = 0.009), circular SD (adjusted p = 0.009), and swing count (adjusted p = 0.024). As in the female analysis, clasping and non-clasping knockout animals did not differ significantly on any of these measures (mean resultant length: adjusted p = 1.000; circular SD: adjusted p = 0.981; swing count: adjusted p = 0.846), demonstrating that the geometric phenotype is consistently present across knockout animals regardless of whether overt clasping occurs during the assay.

It should be noted that the non-clasping subgroup comprised of 6 animals in the case of knockouts and 7 animals in the case of heterozygous and results should therefore be interpreted with appropriate caution given the modest sample size, although the direction and magnitude of effects were consistent across both groups. Together, these findings demonstrate that VECTR-Clasp’s continuous snout-tail trajectory measures capture a genotypic kinematic phenotype that is independent of classical binary clasping classification and detects altered motor dynamics in affected animals that would be classified as phenotype-negative by standard scoring.

### Genotype Classifier Performance

To contextualise the continuous VECTR-Clasp measures against a standard supervised approach, we trained sex-stratified genotype classifiers on the SimBA-extracted pose features. We included these classifiers not as an analytical endpoint but to position VECTR-Clasp relative to a standard supervised approach. The two are complementary rather than competing: a supervised classifier requires genotype labels and at least two groups and returns only a between-group discrimination probability, whereas VECTR-Clasp returns dimensioned, per-animal measures that remain defined for a single animal, a single genotype, or a treatment time-course, settings in which a discrimination probability does not exist.

Sex-stratified random forest genotype classifiers trained on SimBA-extracted pose features achieved above-chance genotype discrimination in both models across 100 repeated balanced-training splits, with a fixed n per group held in for training (n=8 for females, n=10 for males) and remaining animals held out for test. The female wildtype vs. heterozygous classifier achieved a mean accuracy of 77.4% (sensitivity 74.8%; specificity 92.7%), while the male wildtype vs. knockout classifier achieved a mean accuracy of 82.8% (sensitivity 77.1%; specificity 89.8%). The lower accuracy in the female model is consistent with the more subtle phenotype expected in heterozygous versus full knockout animals. These results confirm that DeepLabCut-derived pose features carry meaningful genotypic signal.

### Feature Importance

SHAP analysis on representative median-accuracy splits identified bilateral hind-limb adduction angles as the primary genotype-discriminating feature family in both models (Fig 7). In the male classifier, the single highest-importance feature was the temporal standard deviation of the angle formed between bilateral back paws at the body centre (mean |SHAP| = 0.068), followed by mean bilateral back knee adduction angle (mean |SHAP| = 0.060). In the female classifier, mean bilateral back knee adduction angle was the strongest predictor (mean |SHAP| = 0.043), followed by bilateral back paw angle (mean |SHAP| = 0.041). All 10 features in both models were primarily angular measures between body landmarks (Fig 7 A,B). Although importantly, snout tip vertical movement variability (snout_tip_y_sd) emerged as the 6th most important feature in the female model (mean |SHAP| = 0.016), independently identifying snout movement dynamics as a genotype-discriminating signal which is directly related to VECTR-Clasp’s measurement axis.

**Fig 7.**
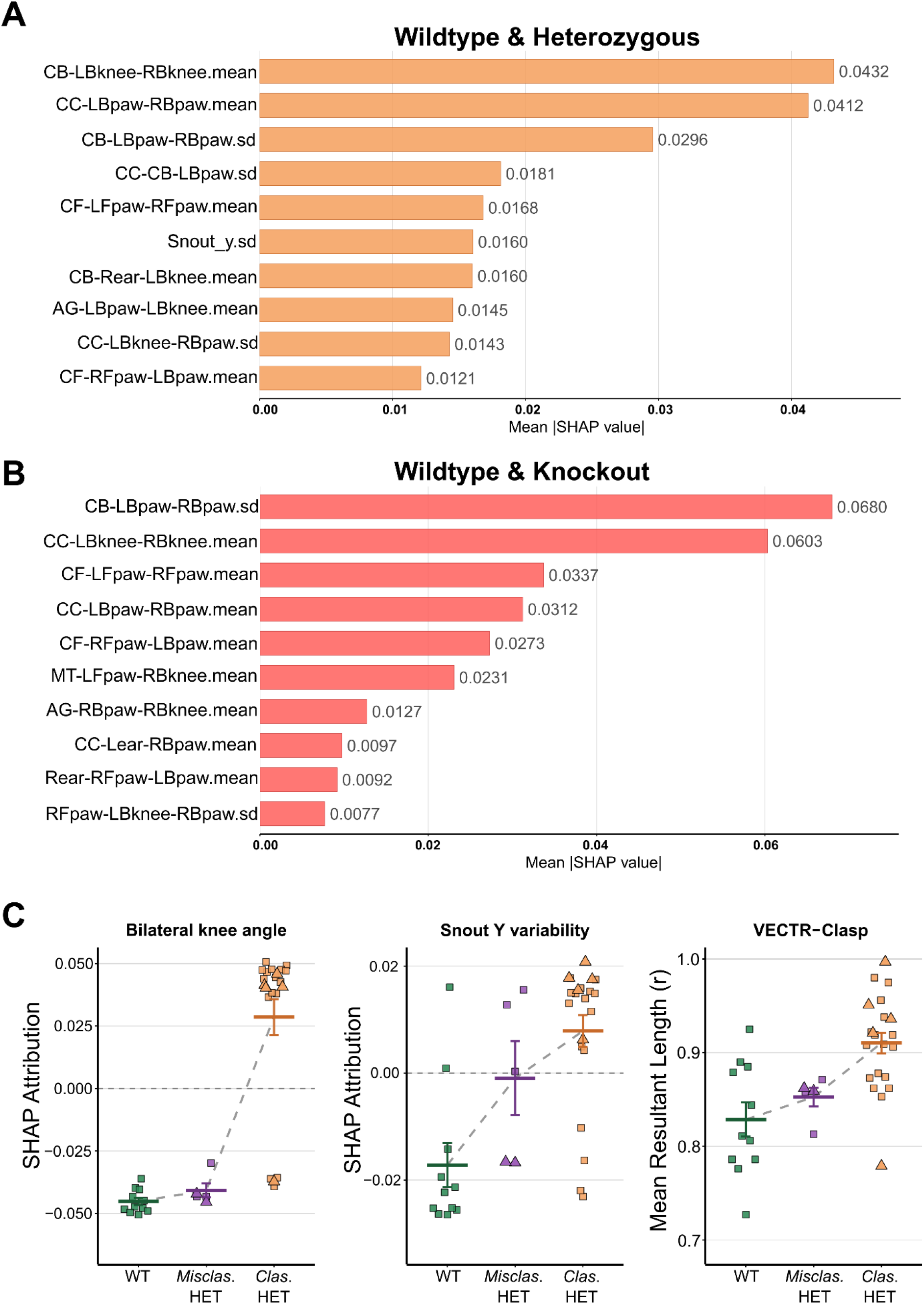
SHAP feature importance and gradient phenotypic signal across classification approaches. (A) Top 10 SHAP feature importances (mean |SHAP value|) from the random forest genotype classifier trained on female wildtype vs heterozygous animals. (B) Top 10 SHAP feature importances from the classifier trained on male wildtype vs knockout animals. (C) Three-panel comparison across wildtype (WT), random forest-misclassified heterozygous (Misclas. HET), and correctly classified heterozygous (Clas. HET) animals. Left: SHAP attribution for the top-ranked bilateral knee angle feature, showing a bimodal distribution in which misclassified animals are indistinguishable from wildtype. Centre: SHAP attribution for snout vertical position variability (6th-ranked feature), showing an emerging partial gradient. Right: VECTR-Clasp mean resultant length, providing the clearest ordered gradient across groups. Individual data points shown; horizontal lines indicate mean ± SEM. Square symbols = clasping; triangle symbols = non-clasping.

### VECTR-Clasp provides a continuous single-animal representation distinct from genotype classification

Five female heterozygous animals were misclassified as wildtype by the genotype classifier on the held-out test set. Examination of their SHAP attribution values for the top discriminating feature (bilateral knee angle) revealed that all five animals showed negative attribution values (mean = −0.041), indistinguishable from the WT group (mean = −0.045) and opposite to the correctly classified heterozygous animals (mean = +0.029). The classifier’s primary postural feature assigned them a wildtype-like signal.

In contrast, examination of the same five animals using VECTR-Clasp continuous measures revealed an ordered intermediate phenotypic position across primary measures: mean resultant length 0.853 (vs. wildtype mean 0.829 and correctly classified heterozygous mean 0.910); circular SD 0.564 (vs. wildtype 0.606 and correct heterozygous 0.415); swing count 46.8 (vs. wildtype 52.8 and correct heterozygous 26.2). This gradient was consistent across all five animals and all three measures. Notably, two of the five misclassified animals showed zero clasping duration, yet their mean resultant length values (0.859 and 0.862 respectively) exceeded the wildtype group mean, demonstrating that VECTR-Clasp detects phenotypic signal in these animals despite their binary classification as phenotype-negative.

Given that this comparison is based on 5 misclassified animals from a single representative holdout split, these findings are illustrative rather than quantitative. Notably, the elevation in mean resultant length was present across all five animals irrespective of their position on the classifier’s continuous decision axis: the two animals assigned confidently wildtype probabilities (P(HET) = 0.20 and 0.21) showed mean resultant length comparable to, or higher than, the three animals nearer the decision boundary (P(HET) = 0.43, 0.44, and 0.49). This indicates that the directional phenotype detected by VECTR-Clasp does not appear to depend on the static posture features prioritised by the classifier, though as noted below the two approaches overlap substantially when assessed across the full held-out set.

A three-panel comparison of SHAP attributions and VECTR-Clasp measures across wildtype, RF-missed heterozygous, and RF-correct heterozygous groups further illustrates the progression from bimodal to gradient signal (Fig 7C): the static bilateral posture feature (bilateral knee SHAP) shows a bimodal pattern with misclassified heterozygous mice indistinguishable from wildtypes; the snout movement variability SHAP feature (6th-ranked) shows an emerging partial gradient (still split by clasping/non-clasping mice); and VECTR-Clasp mean resultant length provides the clearest, fully ordered gradient. This demonstrates that as the measurement approach moves from aggregate static posture toward continuous whole-body movement dynamics, phenotypic signal becomes progressively more recoverable in phenotypically subtle animals.

To assess directly whether VECTR-Clasp captures information beyond the classifier’s continuous output rather than only its discrete label, we related each VECTR-Clasp measure to the classifier’s continuous decision value across the full held-out set. The decision value was moderately-to-strongly correlated with each geometric measure in both models (female wildtype/heterozygous, Spearman ρ: mean resultant length 0.56, circular SD -0.58, swing count -0.55; male wildtype/knockout: 0.58, -0.58, -0.70; all p < 0.05). Partial correlations between genotype and each measure, controlling for the continuous decision value, revealed no residual genotype association in either model (all partial p > 0.30). These analyses are underpowered at the present sample sizes, and their point estimates should be interpreted with caution. For the specific task of discriminating genotypes, VECTR-Clasp and the supervised classifier therefore draw on overlapping variance which is an expected result, since both are computed from the same pose data. This overlap concerns only the between-group discrimination contrast; it does not diminish the distinct purpose of the geometric measures, which is to provide a continuous, model-free description of individual animals in settings where a discriminative probability is undefined.

## Discussion

Behavioural assessments in mice are critical tools for studying brain function and the impact of genetic and other manipulations. Here, we report a pipeline for automated quantitation of clasping behaviour in mice and a pipeline and package to extract continuous kinematic data from the assay. We demonstrate the application of VECTR-Clasp using *Cdkl5-*deficient mice to reveal additional quantitative phenotypes. The findings provide automation of an important assay for CDD research as well as potential applications in the broader area of mouse models of ataxia and neurodevelopment or neurodegenerative disease. The hind-limb clasping assay is widely used in the neuroscience field but remains based on a 5-point scale (11), and it is a commonly used assay in CDKL5 associated research (15,16). To further improve the readouts from this assay, we provide a method through which we can quantify other aspects of this assay to assess angular distribution of snout trajectories and general movement parameters which accurately captures genotype differences in CDD not previously described.

DeepLabCut and SimBA were developed in order to bridge the gap between machine learning and behavioural neuroscience (19,20). Here, we used them together and both provided highly accurate results demonstrating a reproducible pipeline with modest sample sizes. DeepLabCut was successfully used to accurately detect body parts in a 2D projection although a 3D approach may provide further objectivity, particularly in the context of individual limb movement and relation (33). This would account for mice spinning on their tail axis, in turn making accurate measurements of angles and mechanics of hind limbs more feasible. Also, as minor jittering still persisted, more sophisticated filtering approaches could be applied to reduce residual jitter. The SimBA random forest approach provided highly accurate predictions and was able to extract various useful features from the DeepLabCut pipeline. Interestingly, the most important features for prediction were relative geometric positioning of body parts in respect to one another, through using features such as the convex hull perimeter, capturing the geometry of limb retraction.

Although this study focused on the more conservative F0.5 model, an approach using the F1.0 or F1.5 based approach may be more desirable for labs which want to detect all events to maximise false positives and minimise false negatives followed by manually screening for true positives. It is also important to note, that the model was trained by one person meaning that the disparity between the human raters may also be biasing the model as it is learning from one person rather than an intersection vote.

The main known motor phenotypes reported and quantified to date in CDD mice are increased locomotion in home cages and open field (15,34,35), reduced motor coordination in rotarod (15), gait abnormalities (36,37) and displaying hind-limb clasping stereotypies (35). An important finding in the present study was to reveal via analysis of circular statistics and geometric properties previously missed microphenotypes in the exon 6 deletion *Cdkl5* model. This highlights a more nuanced contextual behavioural profile. CDKL5 loss has been largely associated with hyperlocomotion phenotypes in the open field assay in this model (38) while here in this hind-limb clasping assay it is reported that the mice are hypolocomotive in contrast to wildtype counterparts. Notably, there was a reported increase in immobility rates in a 10 minute tail suspension test in an exon 2 deletion model of *Cdkl5* (37). Accordingly, this immobility rate may be linked to the more spatially constrained, less variable pattern of movement during tail suspension of the posture rather than depression-like behaviour. Combined with our study, this indicates that this phenotype of reduced mobility during tail suspension related behavioural assays does appear across different models of *Cdkl5-*deficient mice.

Both heterozygous and knockout animals demonstrated less variable head movement and fewer swinging movement patterns using circular geometric measurements. This indicates that *Cdkl5*-deficient mice show a more spatially constrained movement pattern during this assay than wildtype animals, with lower angular dispersion and fewer swings. We interpret this reduced movement variability cautiously. It may reflect altered motor control, but the present data do not establish which specific motor construct, for example coordination, flexibility, or postural rigidity, underlies it, and we therefore treat these interpretations as hypotheses to be tested rather than established meanings. This is consistent with human patients with CDD (39). It is important to note that this is speculative as usually this mouse model also displays increased locomotion in other assays (15,34,35), combining this assay and analysis technique with another behavioural method such as the open field would provide further interesting information for correlative studies to investigate the head directionality spread across two separate locomotive assays. Notably, we observed differences at baseline between male and female naïve mice with female mice showing more variable movement patterns when compared to males. This highlights that sex is an important consideration when performing this approach.

As pathways such as cerebello-cortico-reticular and cortico-striato-pallido-reticular pathways are implicated in the control of posture during this assay (5), these continuous metrics may allow for more fine-tuned enquiry into these circuits at higher resolution. This is particularly of interest as CDKL5 is expressed abundantly in the striatum (40) and knockout of *Cdkl5* has also been shown to alter inhibitory transmission in the cerebellum (36). This is consistent clinically as MRI studies have shown atrophy of the cerebellum in CDD patients (41). This suggests that these motor pathways may contribute to the reduced movement variability described here. The level of quantification described here could complement electrophysiological or imaging studies aimed at mapping these motor related deficits in the model.

However, the translational relevance of hind-limb clasping as an assay must be interpreted carefully. Mice are quadrupedal and the clasping reflex reflects a species-specific postural response that has no direct human equivalent. Motor deficits in CDD patients manifest primarily through impaired acquisition of bipedal motor milestones such as independent walking and postural control (39). The kinematic features described here, including reduced movement variability and constrained snout trajectories, may reflect underlying deficits in motor circuit function that are conserved across species, but direct correspondence between mouse clasping microphenotypes and human motor outcomes remains to be established. Combining VECTR-Clasp with assays that have clearer translational analogues, such as gait analysis or open field, would strengthen the bridge between preclinical findings and clinical motor endpoints in CDD.

A central motivation for developing continuous kinematic frameworks such as VECTR-Clasp is that traditional categorical scoring systems may lack the sensitivity required to detect subtle treatment effects in preclinical models. The standard manual 5-point clasping scale captures only overt phenotypic severity, meaning that a therapeutic intervention producing a genuine but modest improvement in motor dynamics could go undetected if it does not shift an animal across a categorical threshold. Continuous microphenotype measures, by contrast, are sensitive to gradations of improvement below that threshold, making them potentially more informative for evaluating treatments at early or intermediate stages of efficacy. This is of particular relevance to CDD, where therapeutic strategies including CDKL5 protein replacement and other approaches are under active investigation (17), and where detecting partial rescue of motor function in preclinical models could meaningfully inform clinical translation.

The geometric readouts from the present study may also be useful for other assays used in the preclinical study of neurological disorders. Specifically, we hypothesise that the tail suspension test will be analysable using this framework and would be particularly useful in depression models as the immobility measure is easy to derive if following this pipeline (42). Although, we note that this has not been shown here directly in this study.

Further developments in computational behavioural neuroscience are necessary. As behaviour in its most fundamental form exists as continuous changes in space and time, it can be most naturally expressed as vectors. Such as in the case of place cells (2), head direction cells (43), and grid cells (44), being discovered due to behavioural correlates with neuronal activity, it becomes natural to assume there may be similar cells, but behavioural neuroscience has lacked the granularity or resolution to accurately detect them. Here, it has been demonstrated that open-source tools and frameworks can be utilised to accurately and quantitatively measure behaviours such as the hind-limb clasping assay at a finer resolution which provided a more comprehensive, and richer, phenotyping of motor impairment in *Cdkl5-*deficient mice.

### Limitations of the study

Several limitations of the present study warrant consideration. The DeepLabCut and SimBA classifier components of the pipeline were trained exclusively on adult *Cdkl5*-deficient mice and their wildtype littermates, and while the inclusion of female heterozygous animals in the SimBA validation demonstrates cross-sex applicability of the geometric framework, formal retraining and validation of the behavioural classifier across developmental timepoints remains an important next step. More broadly, the generalisability of VECTR-Clasp to other mouse models of neurological disease in which hind-limb clasping is a core phenotype such as Huntington’s disease (13), cerebellar ataxia (11), and Parkinson’s disease (8) models has not yet been tested. The hind-limb clasping assay is widely used across these models, and the geometric framework operates on standard DeepLabCut outputs, meaning adaptation requires only retraining of the pose estimation model on the target species and recording setup. Nevertheless, whether the specific kinematic microphenotypes described here reflect features unique to CDKL5 loss-of-function or represent a more general signature of motor dysfunction will require cross-model investigation. Additionally, body weight was not directly measured in the present cohort, a proxy body size metric was included as a covariate in linear models, but weight differences between genotypes cannot be fully excluded as a contributing factor to the observed kinematic differences. The pixel-based ratio used here captures relative body proportions but does not account for differences in body mass, fat distribution, or muscle bulk that may independently influence movement dynamics. Future studies should incorporate direct weight measurements at the time of recording to fully exclude this confound. Furthermore, the age covariate analysis was limited to wildtype animals and cannot speak to potential age × genotype interactions in kinematic measures across the window examined here.

A further consideration concerns our decision to quantify clasping as a discrete, full-clasping event rather than as a continuous measurement of individual hind-limb position. During tail suspension, mice frequently rotated about the vertical axis of the suspended tail, and this axial spinning intermittently occluded the hind-limb landmarks in the 2D camera projection. As a consequence, frame-by-frame measures that depend on precise localisation of both hind limbs, such as inter-limb adduction angles or absolute retraction distance, were subject to unstable pose estimation confidence and could not be extracted with the reliability required for quantitative between-group comparison. Rather than report continuous hind-limb metrics of uncertain accuracy, we therefore operationalised clasping as a binary event defined by full retraction of both hind limbs, a criterion that could be scored consistently by both the automated classifier and independent human raters, and which underlies the high inter-rater and human-model agreement reported here. Anchoring the geometric analysis to the more reliably tracked snout and base-of-tail landmarks provided a robust axis for continuous kinematic measurement while avoiding the confound introduced by axial rotation. As noted above, a multi-camera 3D acquisition setup would substantially mitigate this limitation by resolving limb position independently of the animal’s rotation, and represents a natural extension for future work seeking to derive continuous hind-limb kinematics directly from this assay.

Finally, our comparison with a supervised genotype classifier warrants a clear statement of scope. When VECTR-Clasp measures were related to the classifier’s continuous output, they drew on substantially overlapping variance and retained no independent association with genotype after adjustment. We therefore do not claim that the geometric measures recover discriminative information beyond that available to established supervised methods applied to the same pose data; rather, their value lies in providing a continuous, model-free, single-animal representation of movement that is defined where a discriminative classifier is not. Relatedly, although these measures are demonstrably sensitive to genotype, we do not claim that their specific biological interpretation is established, or that it is clearer than that of features surfaced by supervised models. Assigning definitive biological meaning such as for example, distinguishing rigidity, reduced flexibility, or altered coordination, will require convergent evidence from circuit-level manipulation or from assays with clearer translational analogues, and we present the biological framing here as hypothesis-generating rather than settled.

It should also be noted that the genotype classification and feature interpretation approach employed here represents one analytical framework among several viable alternatives. The random forest classifier was selected for consistency with the SimBA pipeline and its established utility in behavioural neuroscience, and SHAP values were used for feature attribution given their interpretability. However, alternative supervised learning approaches such as support vector machines, gradient boosting, or other architectures may offer different performance characteristics depending on sample size and feature structure. Similarly, alternative feature attribution methods such as permutation importance or integrated gradients may recover partially different feature rankings. Additionally, the genotype classifier was underpowered to perform in-depth cross-correlational analysis with VECTR-Clasp. The convergence of SHAP-derived angular features and VECTR-Clasp snout trajectory measures on a consistent genotypic signal suggests the findings are unlikely to be artefacts of the specific analytical choices made here, but systematic comparison across methods remains a worthwhile direction for future work.

## METHODS

### Ethics Statement

All procedures involving animals were performed in compliance with EU Directive 2010/63/EU on the protection of animals used for scientific purposes. Mice were housed at controlled conditions including a room temperature (20°C–25°C), and humidity (40%– 60%), on a 12 h dark-light cycle with ad libitum access to water and food. All procedures were approved by RCSI University of Medicine and Health Sciences’ Research Ethics Committee (REC 1587), and under license from the Ireland Health Products Regulatory Authority (AE19127/P087)

### Animal line

The mouse model of CDKL5 deficiency disorder used was the B6.129(FVB)-*Cdkl5^tm1.1Joez^*/J (15) obtained from the Jackson Laboratory (strain #021967). Offspring heterozygous females were bred in-house with male littermate wildtypes to produce experimental mice. Mice were group-housed, and experiments were performed once mice were P40 or above.

Genotyping was performed using a PCR-based technique to detect the deletion of the *Cdkl5* exon 6 sequence. Using primers with the sequences as follows: Wild type Forward 5’-GGAAGAAATGCCAAATGG AG-3’, Wild type Reverse 5’-GGAGACCTGAAGAGCAAAGG-3’, Mutant Forward 5’-CCCTCTCAGTAAGGCAGCAG-3’, Mutant Reverse 5’-TGGTTTTGAGGTGGTTCACA-3’.

A total of 92 mice were used in this study; 2 mice had insufficient snout and base-of-tail tracking for vector-based analysis and therefore excluded from VECTR-Clasp analysis.

### Hind-Limb Clasping Assay

Mice were suspended by their tail 30cm above the platform, and 30cm from the camera (Logitech C920 HD Pro) for 60 seconds. Video acquisition specifications were 1080x1920 pixels at 30Hz. A total of four recording environments were collected. Calibration was performed with a ruler in the same position to approximate distance on each recording day. All behavioural assays were performed by the same male experimenter each trial. The experimental setup and analysis pipeline are shown in Fig 1.

### Pose Estimation

Frame extraction and labelling was performed with DeepLabCut. Frames were extracted using k-means clustering (80%) and manually chosen clasping frames (20%) ensuring representative postures were selected. 612 frames were manually labelled across 15 videos consisting of 5 wildtype, 6 knockout and 4 heterozygous mice, with 15 body parts annotated (snout_tip, centre_front, centre_centre, centre_back, anogenital_region, base_of_tail, mid_tail, left_ear, right_ear, left_front_paw, right_front_paw, left_back_paw, left_back_knee, right_back_paw, and right_back_knee). Skeleton connections of body parts were created and manually checked to ensure label quality. A 90:10 training-test split was used to generate the dataset, resulting in 551 training frames and 61 test frames overall.

A ResNet-101 backbone with heatmap-based key point detection with location refinement enabled was used. The training settings were as follows: Batch size: 8, Epochs: 200. Optimizer: AdamW, learning rate schedule: 0.0001 until epoch 90, 0.00001 until epoch 120, Seed: 42, Data augmentation: Affine transforms (rotation ±30°, scaling 0.5-1.25, applied with p=0.5). Gaussian noise (σ = 12.75). Motion blur enabled. Cropping: 448×448 px, hybrid sampling, max shift 0.1. Images normalized to RGB. Training was performed with PyTorch using a NVIDIA GeForce RTX 3060 Ti and an Intel i5-12400F.

Once desired evaluation metrics were obtained, the trained model was applied to unseen videos for trajectory extraction. Predictions were filtered using a median filter (5-frame window) to reduce jittering. CSVs then exported for downstream purposes.

### Behavioural Classifier

SimBA generated features from the DeepLabCut CSVs, including raw x-y position values, per-body-part movement metrics, rolling-window statistical summaries, pairwise distances, and body-part angles. These features were used as input for Random Forest classification of clasping behaviour. Five mice (2 wildtype, 3 knockout) were manually annotated in SimBA for the presence or absence of hind-limb clasping bouts. Heterozygous animals were excluded from SimBA classifier training as the aim was to train the model to recognise the binary presence or absence of hind-limb clasping behaviour, rather than genotypic differences in its expression. As clasping represents a stereotyped motor pattern whose topography is consistent across genotypes, wildtype and knockout animals provided the clearest examples of behaviour absence and presence respectively, maximising classifier sensitivity to the behaviour itself independent of genotype.

Although five videos were used for annotation, this yielded 1,772 labelled frame-level samples for classifier training, which was considered sufficient given the constrained and stereotyped nature of the clasping behaviour and the geometric features extracted by SimBA. Classifier generalisation was evaluated on 70 independent held-out animals not used at any stage of training or annotation. The Random Forest classifier was trained on these 1,772 labelled samples, comprising 622 clasping samples and 1,150 non-clasping samples (≈35% vs 65%) across 9000 frames (1800 frames per video) obtained from the 5 mice. The following hyperparameters were used: Number of trees: 2000, Maximum features: sqrt, Criterion: Gini impurity, Minimum samples split: 2, Minimum samples per leaf: 1, Maximum depth: None (unlimited). The training dataset was balanced using SMOTE (Synthetic Minority Oversampling Technique) to account for class imbalance between clasping and non-clasping frames. Classifier performance was evaluated using a confusion matrix, and precision-recall metrics. Discrimination thresholds were chosen based on highest F1.0 and F0.5 values. Clasping events under 400ms were discarded to reduce false positives.

### Human-Machine Agreement

Two blinded experimenters scored 70 independent videos not used at any stage of training (consisting of 27 wildtype, 26 heterozygous, and 17 knockout mice) manually with a binary clasp or no clasp scoring at each second of recording. A clasping event was defined as full retraction of both hind limbs to the centre of the mouse’s body.

The same 70 held-out videos were then analysed using the DeepLabCut-SimBA pipeline to predict clasping bouts. In order to equalise the scoring time resolution, whenever the model predicted a clasping event to occur at any duration in a one second epoch - it was counted as one second. With this, we assess the model’s accuracy at a rate of 1 second resolution.

To provide a realistic ceiling for accuracy and to contrast with human-to-human variability: the human intersection was compared to the F0.5 SimBA model, as a more conservative approach, the human intersection served as the “ground truth”. Cohen’s kappa was then calculated for inter-rater reliability, along with the per second percent agreement overall. All downstream analysis were performed using the F0.5 model.

### Genotype Difference

Clasping analysis was sex stratified and compared across male (wildtype and knockout) and female (wildtype and heterozygous) genotypes. The results were calculated for both human and SimBA model based clasping measurements. To determine if genetically altered mice display clasping versus wildtype counter parts, Fisher’s exact test was performed on mice to detect any mice which displayed clasping. Mice which were predicted to have less than 1 second clasping and present with only 1 bout were manually investigated to confirm or reject the mouse as a clasper.

### Vector Based Geometry

To extract further information from the videos a vector based geometric approach was performed focusing on snout trajectory as a pattern of movement during tail suspension. Videos were assessed to count how many frames had a likelihood of above 0.6 in both snout and base-of-tail, videos with less than 50% of frames present were discarded.

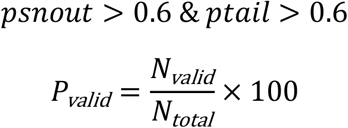

#### Relative Snout Tip Coordinates

To analyse the snout’s movement relative to the body, the base of the tail was treated as the origin (0,0). The snout tip’s coordinates were then recalculated relative to this anchor point.

Let:

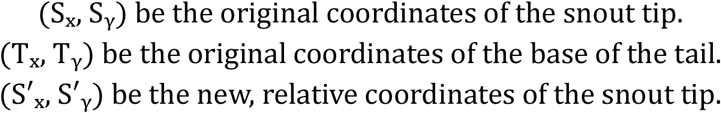

The formula for the relative coordinates is a simple vector subtraction (Equation 1):

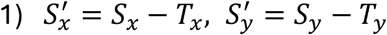

#### Polar Coordinates (Radius and Angle)

To quantify the swinging motions, the relative Cartesian coordinates were converted into polar coordinates.

The radius (*r*) represented the distance from the base of the tail to the snout tip. It was calculated using the Euclidean distance formula (Equation 2).

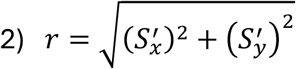

The angle (θ) represented the direction of the snout tip relative to the base of the tail. It was calculated using the atan2 function to ensure the angle is correct across all four quadrants. The corresponding result was in radians (Equation 3).

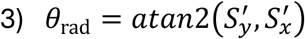

To convert the angle from radians to degrees Equation 4 was used:

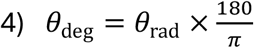

#### Head Movement (Frame-to-Frame Distance)

To quantify the overall activity level, the Euclidean distance the snout tip travels between consecutive frames was calculated (Equation 5a & 5b).

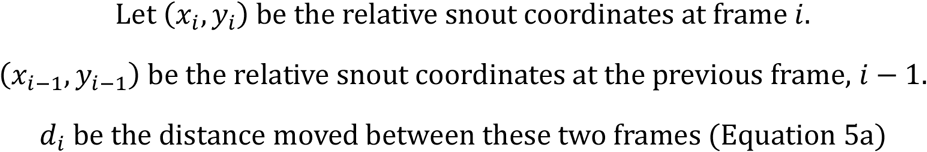

The formula is:

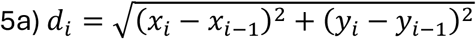

The Total Distance is the sum of these movements over all frames (Equation 5b):

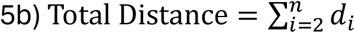

#### Circular Statistics

Circular statistics were then used to quantify directional data.

The mean resultant length measures the concentration of the angles (Equation 6). It ranges from 0 (angles are uniformly distributed) to 1 (all angles are identical). A higher value indicates less angular variation.

For a set of n angles:

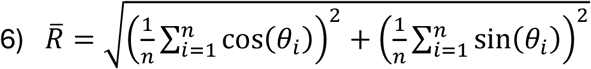

The Circular Standard Deviation (σ) measures the dispersion of angles around the mean angle. It is derived from the mean resultant length (Equation 7). The result is given in radians.

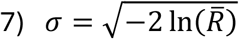

#### Swing Count Calculation

This was not a single formula but an algorithm to count the number of swings across a central line. Each frame’s angle (θ) is categorized relative to a midline (90^∘^) and a deadband (45^∘^) which was halfway between the hanging position and the side of the tail base on both sides. An animal was deemed to be swinging to the left if θ > (midline plus dead band), swinging to right if θ < (midline minus dead band), while in the centre as within the dead band zone.

### Genotype Classifier and SHAP Analysis

To contextualise VECTR-Clasp’s contribution relative to standard supervised classification approaches, two sex-stratified binary random forest genotype classifiers were trained on DeepLabCut derived pose features extracted from SimBA (total features: 2739). One model compared male wildtype versus knockout and one comparing female wildtype versus heterozygous, both reflecting clinically relevant comparisons. SimBA’s feature extraction pipeline was applied per-animal to generate a feature matrix comprising per-video mean and standard deviation statistics for landmark angles and raw coordinate positions. Two male animals were excluded from VECTR-Clasp geometric analysis because fewer than 50% of frames met the snout and base-of-tail likelihood threshold (>0.6) required for vector-based analysis. These animals were retained for the genotype classifier, which uses the broader SimBA feature set that does not depend specifically on snout-tail tracking, bringing the total to 77 videos outlined in Table 1. Features with near-zero variance were removed, the top 300 features by variance were retained for modelling followed by removal of features with pairwise Pearson correlation exceeding 0.90 to reduce redundancy.

Each classifier was implemented as a random forest (number of trees = 1000) and was evaluated across 100 repeated balanced-training splits, with a fixed n per group held in for training (n=8 for females, n=10 for males) and remaining animals held out for test. Per-split accuracy, sensitivity, and specificity were recorded. The representative median-accuracy split was identified and used for SHAP-based feature interpretation. For the female wildtype vs. heterozygous model, the representative median-accuracy split comprised 3 wildtype and 18 heterozygous animals in the held-out test set; of these, 5 heterozygous animals (28%) were misclassified as wildtype and form the basis of the VECTR-Clasp gradient comparison in Fig 7C. For the male wildtype vs. knockout model, the representative split comprised 9 wildtype and 11 knockout animals in the held-out test set, with 1 knockout animal misclassified as wildtype, consistent with the stronger and more penetrant phenotype in full knockout animals.

SHAP (SHapley Additive exPlanations) attribution values were computed applied to the random forest trained on the representative split, with attributions computed across all animals (train and test combined). Mean absolute SHAP value per feature was used to rank feature importance. To directly assess whether VECTR-Clasp recovers phenotypic signal that binary classification cannot, VECTR-Clasp measures (mean resultant length, circular SD, swing count) were examined in female heterozygous animals misclassified as wildtype by the genotype classifier on the held-out test set. These animals were compared against wildtype animals and correctly classified heterozygous animals using the same measures.

To evaluate whether VECTR-Clasp measures carry genotype information beyond the classifier’s continuous output, the random forest class probability from the representative median-accuracy split was correlated (Spearman) with each VECTR-Clasp measure across the full held-out set for each model. Partial correlations between genotype and each measure were computed using Pearson partial correlation. Given the modest sample sizes (female n = 21; male n = 15), these analyses were treated as descriptive rather than as formal hypothesis tests.

### Statistics

Pose estimation was assessed using values of: Training/validation loss, Root Mean Square Error (RMSE), RMSE at p-cutoff 0.6, Mean Average Precision (mAP) and a Mean Average Recall (mAR).

Behavioural classifier training was assessed using confusion matrices (true positives, true negatives, false positives, false negatives) and Precision-Recall curves. Discrimination Thresholds were determined using the maximum value obtained for F1.0 and F0.5. Human inter-rater variability and agreement was calculated using Cohens Kappa and percentage agreement per second across all 70 independent videos and summated to overall representations. The F0.5 model was assessed similarly to the human intersection to determine accuracy of the model.

A Spearman Correlation was performed to assess the accuracy of duration of time spent clasping reported by the human intersection score and the model score with both R and *p* value reported as metrics of accuracy and significance.

Comparisons use per-animal circular summary statistics assessed between groups. Normality was assessed by Shapiro-Wilk. Subsequently, non-normally distributed data was assessed by Mann-Whitney U (graphs are displayed as the median plus IQR) while normally distributed data was assessed using an unpaired T-Test with Welch’s correction (graph displayed as mean plus standard error of the mean (SEM)). To account for multiple testing in the vector-based geometry assessment, the Bonferroni correction was performed to adjust p values. Significance was assessed at alpha level 0.05.

To test whether VECTR-Clasp measures reflect a genotypic phenotype independent of overt clasping behaviour, animals within each affected genotype were stratified post-hoc into two subgroups: those exhibiting zero clasping duration by both human and automated scoring and those with at least one clasping event. Each VECTR-Clasp measure was then compared across three groups per sex, using one-way ANOVA with Tukey HSD post-hoc correction where data were normally distributed, or Kruskal-Wallis with Dunn’s post-hoc test (Bonferroni correction) where normality was violated.

To assess the effects of sex, age, and body size on continuous VECTR-Clasp outcome measures, linear models were fitted in R. As the gene of interest is X-linked, genotype and sex are structurally confounded in this experimental design. Accordingly, we performed analysis on wildtype male and females to determine if sex alone is a predictor in this analysis method. Age (postnatal days) and a continuous body size metric (tail base to snout: waistline length in pixel, ratio) were included as covariates.

All machine learning pipelines were conducted in a Conda (version 25.5.1) environment with Python version 3.10.18 for DeepLabCut (version 3.0.0rc10) and Python version 3.6.10 for simba-uw-tf-dev (version 3.8.8) while statistics and graphical generation were performed using R Studio (version 2025.9.0.387).

VECTR-Clasp is downloadable as a complete package on GitHub at https://github.com/Jordan-Higgins/VECTR-Clasp, along with example DeepLabCut output CSVs of 6 wildtype and 6 knockout mice during hind-limb clasping and an explanatory guide detailing commands usable with the package.

## Supporting information

Supplementary Video 1 - Wildtype

Supplementary Video 2 - Knockout

## Data and code availability

All original code has been deposited under the MIT licence at the following GitHub repository and is publicly available at: https://github.com/Jordan-Higgins/VECTR-Clasp alongside example DeepLabCut output data for wildtype and knockout mice and simulated data with known results for testing of installation. Raw pose estimation files and summarised behavioural data needed to reproduce all analysis are provided in Supporting Information 1.

## Acknowledgements

The authors would like to thank Dr. Amaya Sanz Rodriguez, and the FutureNeuro operations team for their valuable support throughout this work. The authors have reviewed and edited the output and take full responsibility for the content of this publication.

## Author Contributions

Original conception of the work: J.H, O.M. Contribution and design of the experiment: J.H, O.M. Collection of data: J.H. Analysis: J.H., S.E, K.H, B.E-M, O.M. Data Interpretation: J.H, D.C.H, O.M. Drafting the article or revising it for important intellectual content: J.H., D.C.H, O.M. All authors approved the final version of the manuscript.

## Supplementary Videos

Supplementary Video 1: Representative 60-second tail suspension recording of a wildtype mouse, illustrating the dynamic snout movement and lateral swinging behaviour characteristic of wildtype animals during the assay.

Supplementary Video 2: Representative 60-second tail suspension recording of a *Cdkl5* knockout mouse that did not display overt hind-limb clasping during the trial, illustrating the constrained snout trajectory and reduced lateral movement detected by VECTR-Clasp geometric analysis in the absence of classifiable clasping behaviour.

## Supplementary Data

**Supplementary Fig 1:**
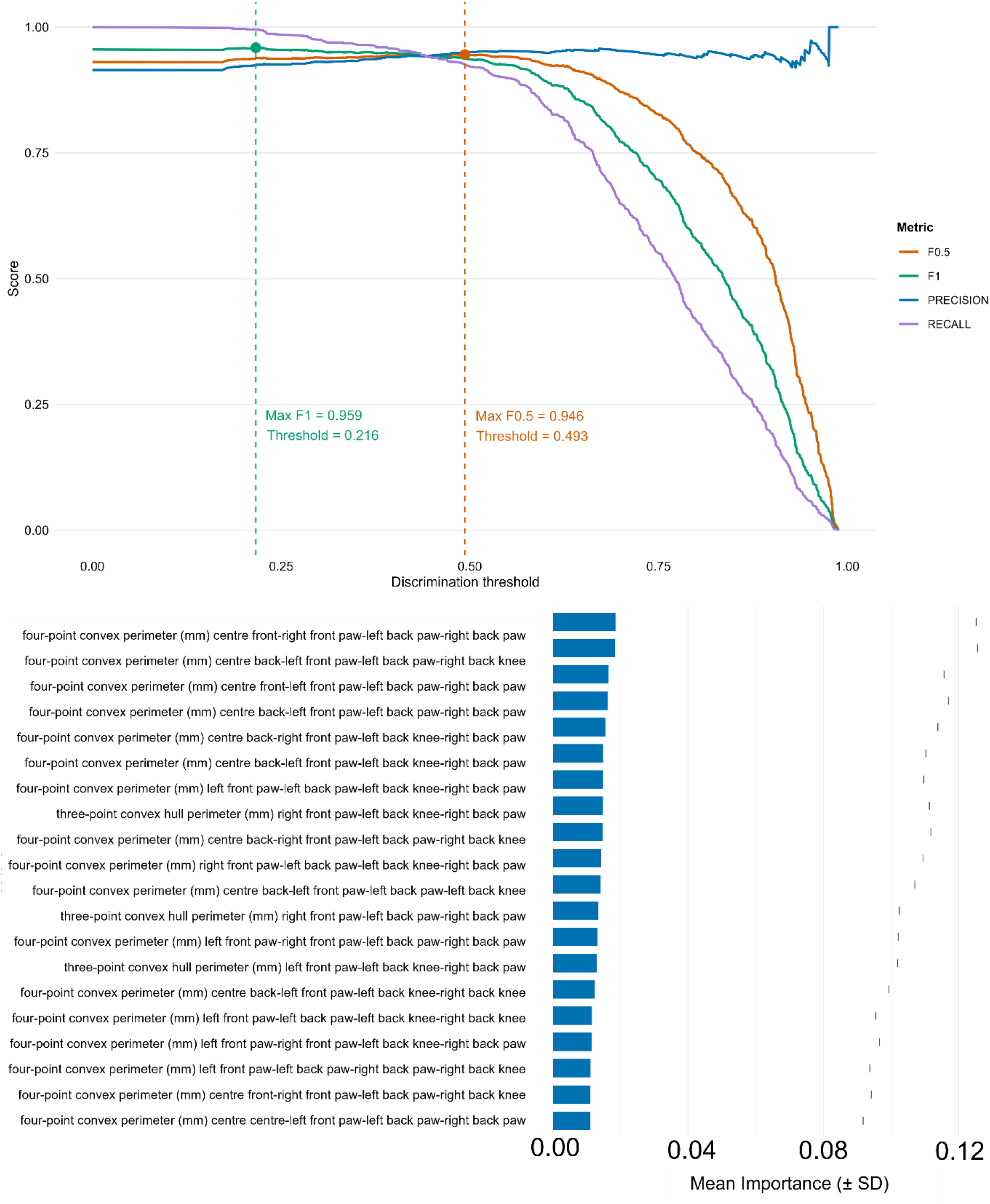
Model Optimization and Feature Importance in Automated Clasping Behaviour Classification. (Top) Precision-recall analysis illustrating the relationship between discrimination threshold and performance metrics. Precision (blue), recall (purple), F1.0 (green), and F0.5 (orange) scores are plotted across thresholds. Dashed vertical lines indicate the discrimination thresholds corresponding to the maximum F1.0 (0.959 at threshold = 0.216) and F0.5 (0.946 at threshold = 0.493). (Bottom) Top 20 most important features contributing to clasping behaviour classification, shown as mean importance ± SD.

**Supplementary Fig 2:**
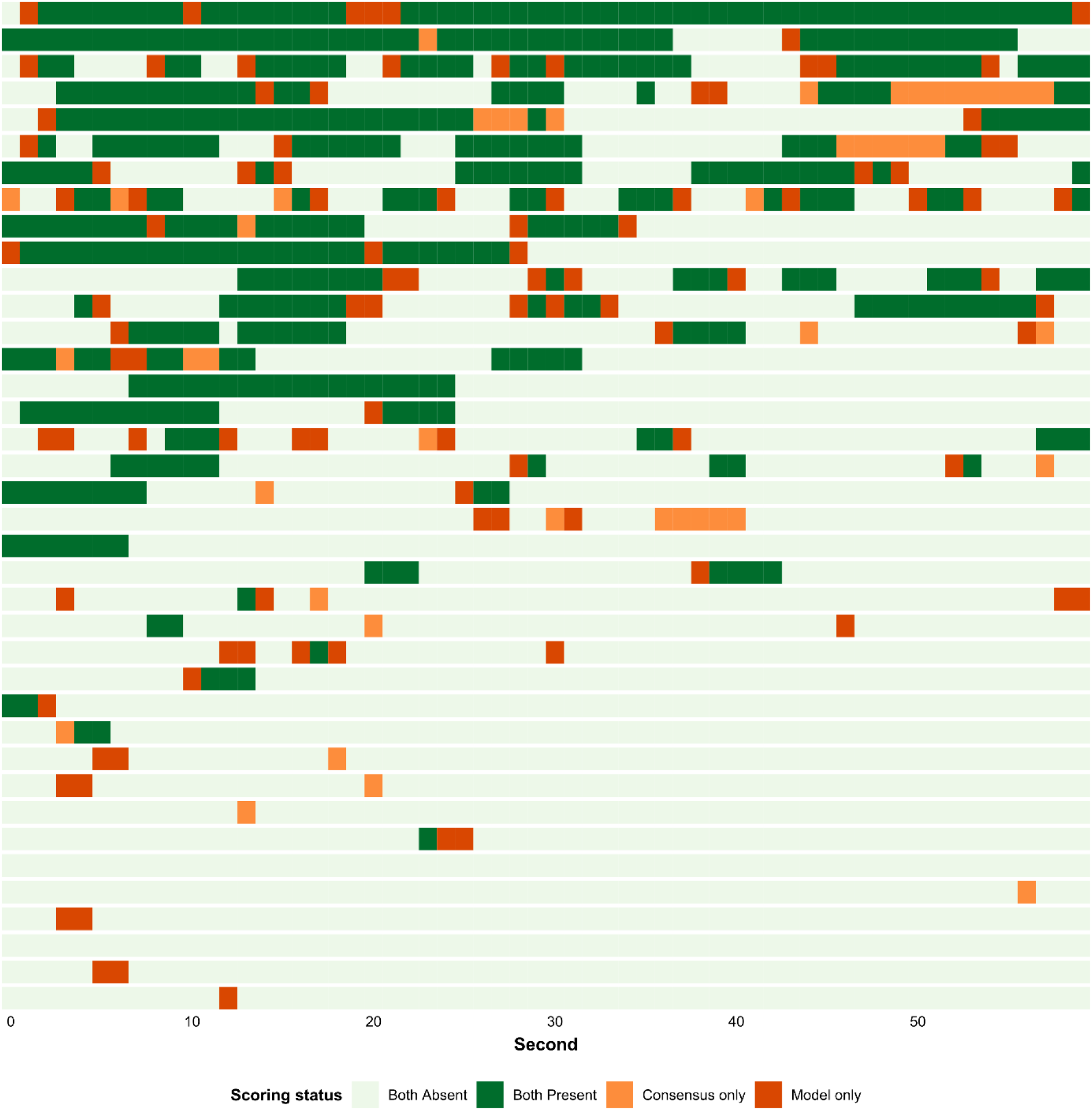
Temporal agreement between automated model scoring and human consensus across individual animals. Each row represents an animal from the held-out SimBA validation set including all animals with at least one bout, with time displayed in one-second epochs across the 60-second trial on the x-axis. Scoring status at each second is colour-coded according to agreement between the SimBA model and the human consensus label (intersection of two independent raters): both absent (light green, no clasping detected by either scorer), both present (dark green, clasping detected by both), consensus only (orange, clasping detected by human consensus but not the model; false negative), and model only (dark orange/red, clasping detected by the model but not the human consensus; false positive). The predominance of light green across the raster reflects the high overall per-second agreement (96.4%), with disagreements largely around the onset and offset of clasping bouts, consistent with the known difficulty of defining precise behavioural boundaries at one-second resolution for both human raters and the automated classifier.

**Supplementary Fig 3:**
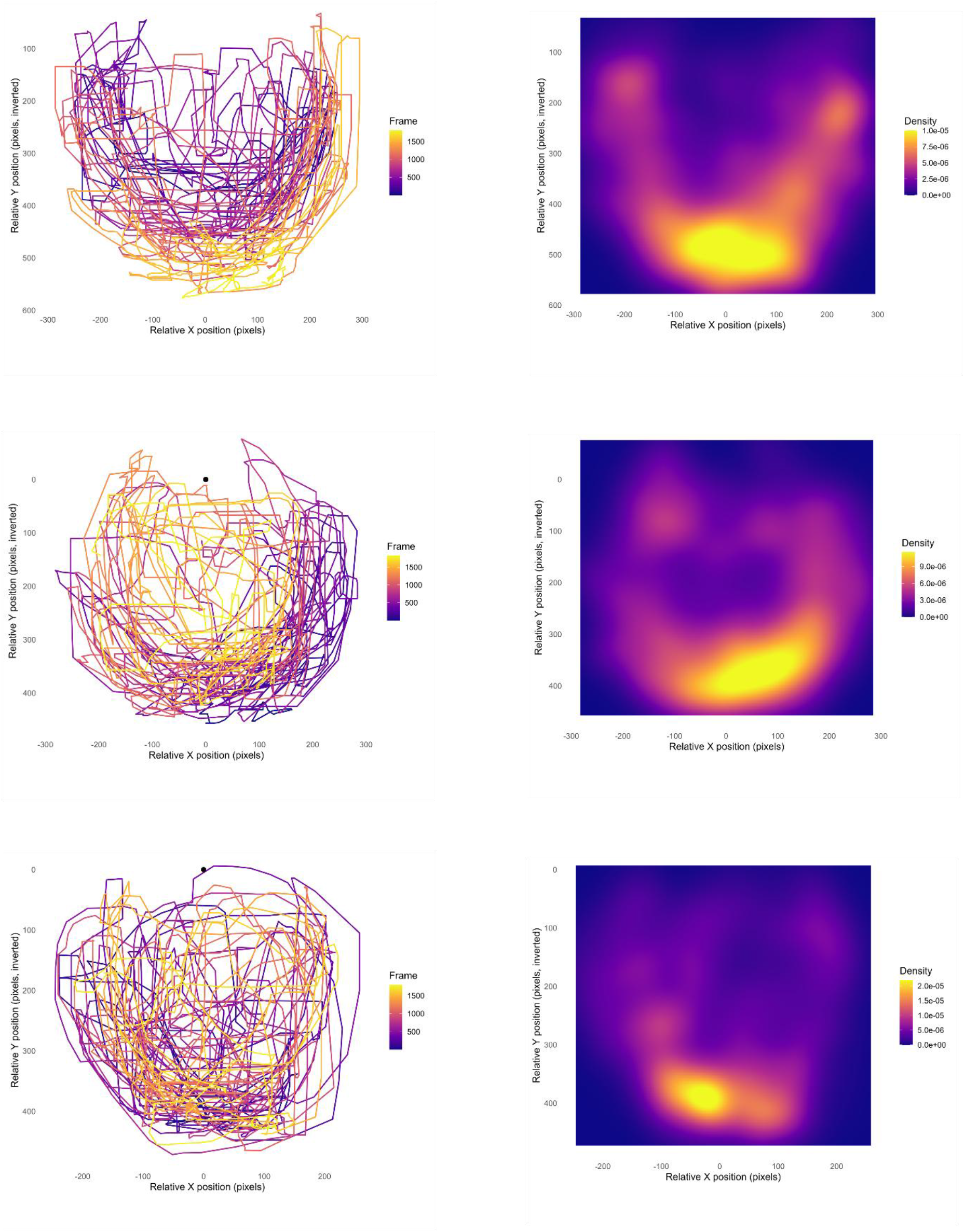
Representative locomotor snout track plots and snout probability density heatmaps for wildtype animals. Each row shows data from an individual wildtype replicate (n = 3). Left column: Two-dimensional track plots of relative position (in pixels) over the recording period. Line colour encodes frame number (purple to yellow, represents early to late), illustrating the temporal progression of movement. Right column: Snout density estimates of position occupancy across the entire recording, with warmer colours (yellow) indicating regions of higher probability density and cooler colours (purple) indicating lower density.

**Supplementary Fig 4:**
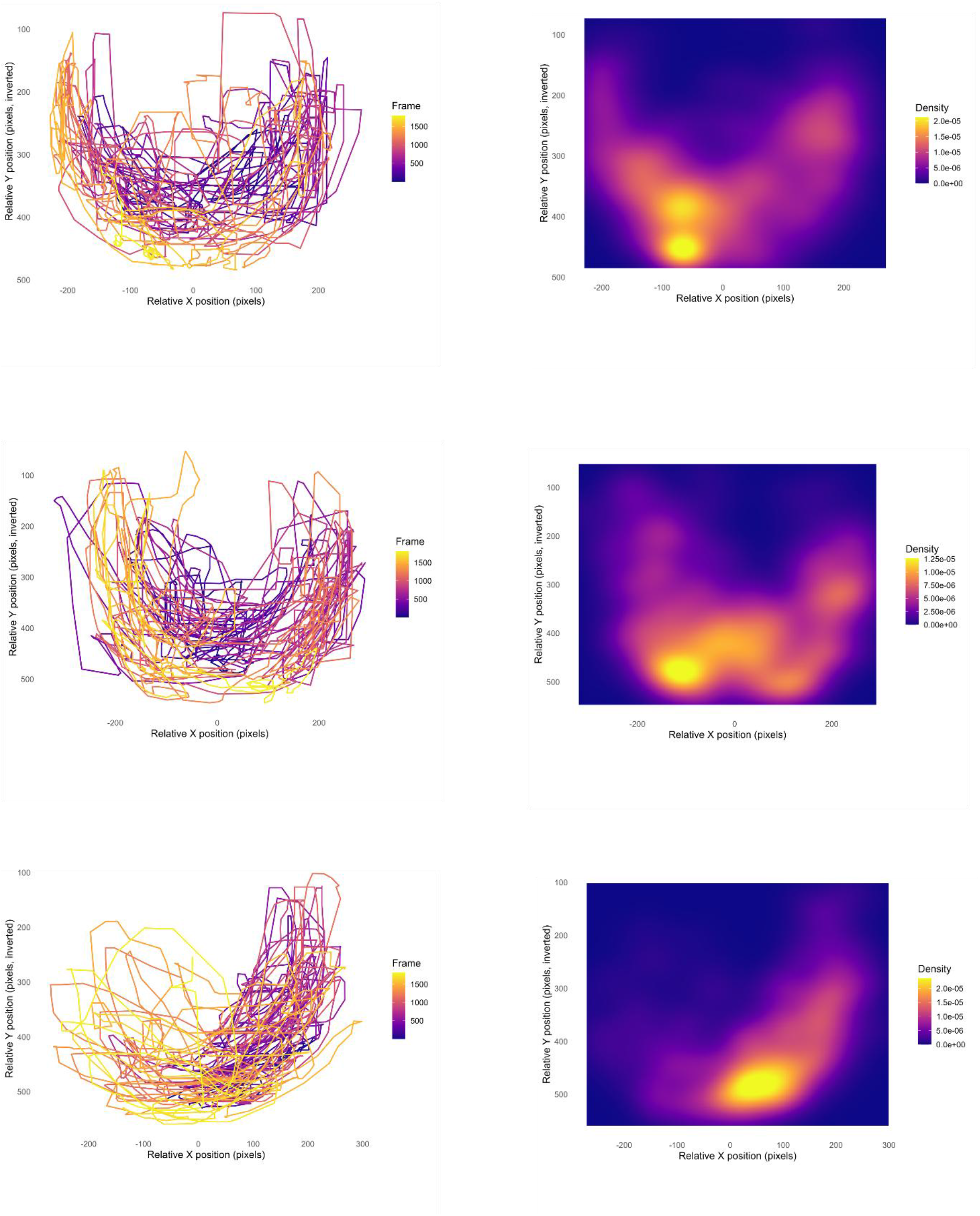
Representative locomotor snout track plots and snout probability density heatmaps for heterozygous animals. Each row shows data from an individual replicate (n = 3). Left column: Two-dimensional track plots of relative position (in pixels) over the recording period. Line colour encodes frame number (purple to yellow, represents early to late), illustrating the temporal progression of movement. Right column: Snout density estimates of position occupancy across the entire recording, with warmer colours (yellow) indicating regions of higher probability density and cooler colours (purple) indicating lower density.

**Supplementary Fig 5:**
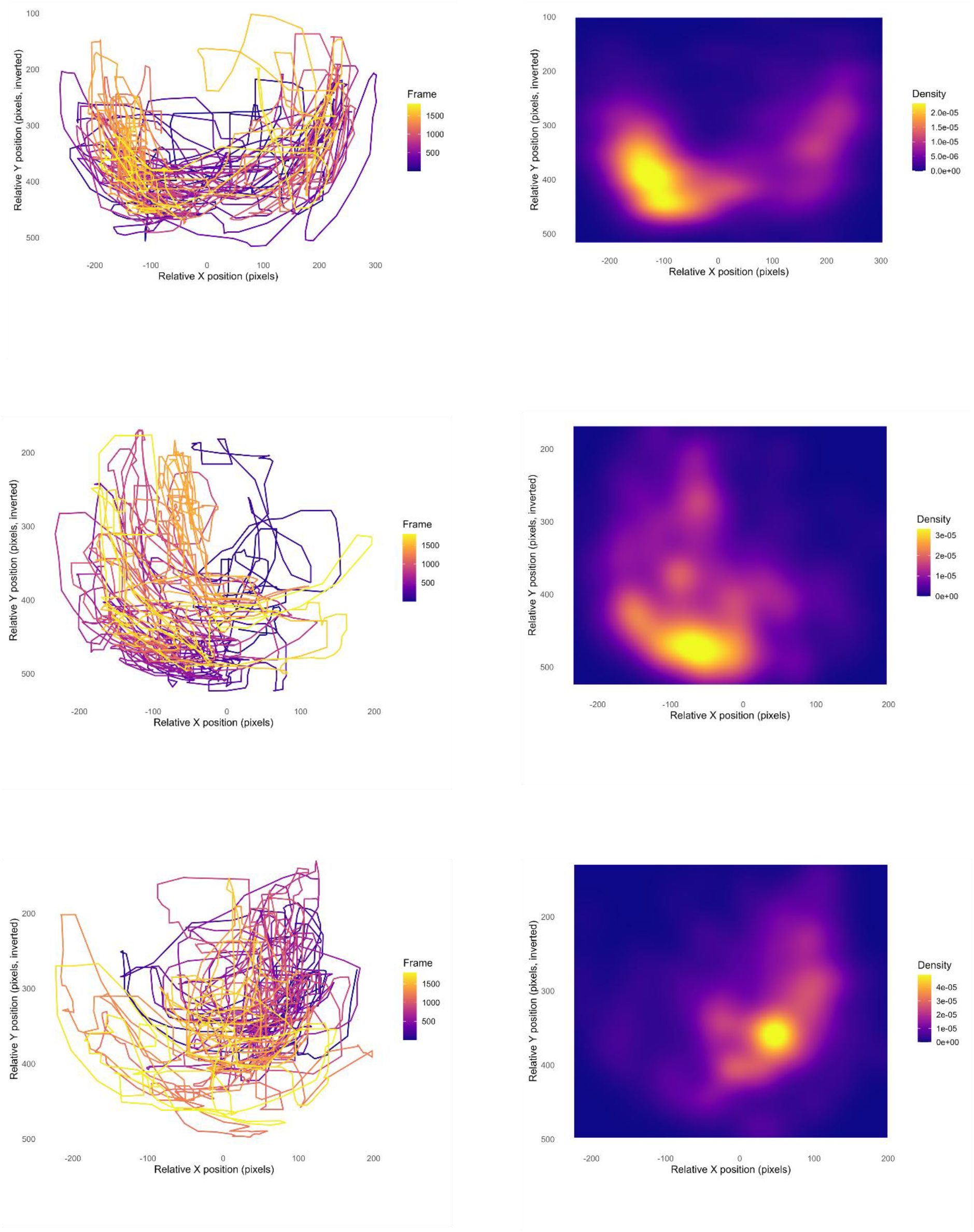
Representative locomotor snout track plots and snout probability density heatmaps for knockout animals. Each row shows data from an individual replicate (n = 3). Left column: Two-dimensional track plots of relative position (in pixels) over the recording period. Line colour encodes frame number (purple to yellow, represents early to late), illustrating the temporal progression of movement. Right column: Snout density estimates of position occupancy across the entire recording, with warmer colours (yellow) indicating regions of higher probability density and cooler colours (purple) indicating lower density.

